# ER-lysosome lipid transfer protein VPS13C/PARK23 prevents aberrant mtDNA-dependent STING signaling

**DOI:** 10.1101/2021.06.08.447593

**Authors:** William Hancock-Cerutti, Zheng Wu, Arun Tharkeshwar, Shawn M. Ferguson, Gerald S. Shadel, Pietro De Camilli

**Affiliations:** Department of Neuroscience, Yale University School of Medicine, New Haven, CT, USA; Department of Cell Biology, Yale University School of Medicine, New Haven, CT; Interdepartmental Neuroscience Program, Yale School of Medicine, New Haven, CT, USA; MD/PhD Program, Yale School of Medicine, New Haven, CT, USA; Howard Hughes Medical Institute; Department of Genetics, Yale School of Medicine, New Haven, CT, USA; Salk Institute for Biological Studies, La Jolla, CA, USA; Program in Cellular Neuroscience, Neurodegeneration and Repair, Yale University School of Medicine, New Haven, CT; Kavli Institute for Neuroscience, Yale University School of Medicine, New Haven, CT; Aligning Science Across Parkinson’s (ASAP) Collaborative Research Network, Chevy Chase, MD

## Abstract

Mutations in VPS13C cause early onset, autosomal recessive Parkinson’s Disease (PD). We have established that VPS13C encodes a lipid transfer protein localized to contact sites between the endoplasmic reticulum (ER) and late endosomes/lysosomes. In the current study, we demonstrate that depleting VPS13C in HeLa cells causes an accumulation of lysosomes with an altered lipid profile, including an accumulation of di-22:6-BMP, a biomarker of the PD-associated leucine-rich repeat kinase 2 (LRRK2) G2019S mutation. In addition, the DNA-sensing cGAS/STING pathway, which was recently implicated in PD pathogenesis, is activated in these cells. This activation results from a combination of elevated mitochondrial DNA in the cytosol and a defect in the degradation of activated STING, a lysosome-dependent process. These results suggest a link between ER-lysosome lipid transfer and innate immune activation and place VPS13C in pathways relevant to PD pathogenesis.

## INTRODUCTION

Genetic studies have revealed many genes whose mutations cause or increase the risk of Parkinson’s Disease (PD). Elucidating the functions of these genes, and the mechanisms by which their mutations cause PD, may provide insights into general PD pathophysiology and yield new therapeutic strategies. Several of these genes have been implicated in mitochondrial function (Malpartida et al., 2021), while many others play a role in the endolysosomal system (Abeliovich and Gitler, 2016; Vidyadhara et al., 2019). The extent of physiological and pathological crosstalk between these two organelle systems is a topic of increasing interest (Hughes et al., 2020; Kim et al., 2021; Yambire et al., 2019).

One of the genes whose mutations are responsible for familial early onset PD is VPS13C (Darvish et al., 2018; Lesage et al., 2016; Schormair et al., 2018). The VPS13C locus was also identified in multiple PD GWAS (Nalls et al., 2019). Additionally, loss-of-function mutations in VPS13C genes were detected in dementia with Lewy Bodies, peculiar protein aggregates enriched in alpha-synuclein and characteristic of PD(Smolders et al., 2021). The VPS13 gene family encodes lipid transfer proteins that localize to a variety of distinct contact sites between membranous organelles(Ugur et al., 2020). These proteins are thought to function as bridges that allow phospholipids to traverse the aqueous cytosolic environment between bilayers through a hydrophobic groove that runs along their length(Guillen-Samander et al., 2021; Kumar et al., 2018; Li et al., 2020).

Initial studies of VPS13C focused on a potential role in mitochondrial physiology (Lesage et al., 2016), as at the time studies of the single yeast Vps13 protein had suggested a role of this protein in the transport of lipids to these organelles(Lang et al., 2015). Such studies reported the presence of VPS13C in mitochondria-associated membrane fractions and showed that VPS13C knockdown causes mitochondrial dysfunction (Lesage et al., 2016). This seemed to be consistent with evidence for a major role of defects in mitochondrial clearance in some familial forms of PD (Pickrell and Youle, 2015). However, subsequent studies showed that while VPS13A (Kumar et al., 2018; Yeshaw et al., 2019) and VPS13D(Guillen-Samander et al., 2021) localize to contact sites between the ER and mitochondria, VPS13C localizes instead to contact sites between the ER and late endosomes/lysosomes (Kumar et al., 2018). The localization of VPS13C at endolysosomes, but not mitochondria, was also supported by proximity labeling experiments(Go et al., 2021; Liu et al., 2018)although genetic evidence for an impact of VPS13C on lysosome properties is still missing. A role of VPS13C in the endolysosomal system is not at odds with the established genetic links to PD, given that many PD genes that function in the endolysosomal system(Abeliovich and Gitler, 2016). How defects in this system promote PD, however, remains to be understood.

Recent studies have implicated activation of the innate immune response in PD pathogenesis. Specifically, it has been reported that defective mitochondrial clearance in mice with Parkin and PINK1 loss-of-function mutations subjected to additional mitochondrial stressors may result in mitochondrial DNA (mtDNA) leakage into the cytosol, leading to activation of the cGAS-STING pathway (Sliter et al., 2018). Such activation, in turn, induces the transcription of interferon-stimulated genes (ISGs) and an NF-кB-mediated inflammatory response (Motwani et al., 2019). While cGAS and STING are primarily expressed in non-neuronal cells in brain, including microglia and astrocytes (Saunders et al., 2018), there is growing evidence that some other genes involved in neurodegenerative diseases, including PD, are also expressed primarily in non-neuronal cells including immune cells(Cook et al., 2017). Interestingly, activation of the cGAS-STING pathway due to elevated cytosolic mtDNA was observed in bone marrow-derived macrophages (BMDMs) and mouse embryonic fibroblasts (MEFs) lacking LRRK2(Weindel et al., 2020), a PD gene associated with lysosomes (Bonet-Ponce et al., 2020). Moreover, absence of C9orf72, an ALS gene and a component of a signaling complex associated with lysosomes (Amick et al., 2016), results in hyperresponsiveness to activators of STING likely due to impaired degradation of STING in lysosomes (McCauley et al., 2020). As multiple groups have shown that activation of STING, an ER resident protein, triggers its transport from the ER to lysosomes, where it is degraded, defective lysosomal function may delay clearance of activated STING (Gonugunta et al., 2017; Gui et al.). These previous studies raise the possibility that an interplay of mitochondrial defects (such as mtDNA leakage) and lysosomal defects (such as impaired STING degradation) may synergize in the activation of the innate immune response leading to neuroinflammation in some neurodegenerative diseases.

Intriguingly, the single yeast VPS13 gene is required both for mitochondrial integrity(Lang et al., 2015; Park et al., 2016) and for proper function of the endolysosomal system(Brickner and Fuller, 1997). Moreover, this yeast protein was identified in a genetic screen for mutations that cause the escape of mtDNA to the nucleus, hence its alias YME3 (Yeast Mitochondrial Escape)(Thorsness and Fox, 1993). Follow-up studies of another YME gene, the ATP-dependent mitochondrial metalloprotease YME1, revealed that escape of mtDNA required degradation of mitochondrial compartments in the vacuole, the yeast equivalent of the lysosome(Campbell and Thorsness, 1998). Moreover, recent studies of mammalian YME1L demonstrate that loss-of-function results in mtDNA leakage and activation of the cGAS-STING pathway (Sprenger et al., 2021). All these findings prompted us to explore a potential activation of the cGAS-STING pathway by mtDNA in VPS13C depleted cells and its potential relationship to lysosome dysfunction.

## RESULTS

### Loss of VPS13C results in perturbation of lysosomal homeostasis

We have previously shown that VPS13C localizes to contact sites between the ER and late endosomes/lysosomes positive for the GTPase Rab7(Kumar et al., 2018), a marker of these organelles(Gillingham et al., 2014). Moreover, VPS13C was a hit in a high-throughput screen for interactors of Rab7a(Gillingham et al., 2019). We have not only confirmed that VPS13C localizes to organelles positive for overexpressed Rab7a or constitutively active Rab7a^Q67L^, but have now demonstrated that co-expression of VPS13C with a dominant negative mutant Rab7a (Rab7a^T22N^), which cannot localize to late endosomes/lysosomes, causes VPS13C to have a diffuse cytosolic distribution (Figure 1A). These findings demonstrate a key role of Rab7 in the recruitment of VPS13C to late endosomes/lysosomes. In this experiment, dispersal of VPS13C to the cytosol, rather than its accumulation in the ER, the other major VPS13C organelle, may reflect insufficient levels of VAP, its ER binding partner.(Kumar et al., 2018) As these experiments corroborate the idea that contacts between the ER and endolysosomes are a main site of action of VPS13C, we investigated whether the absence of VPS13C has an impact on lysosomal parameters.

**Figure 1.**
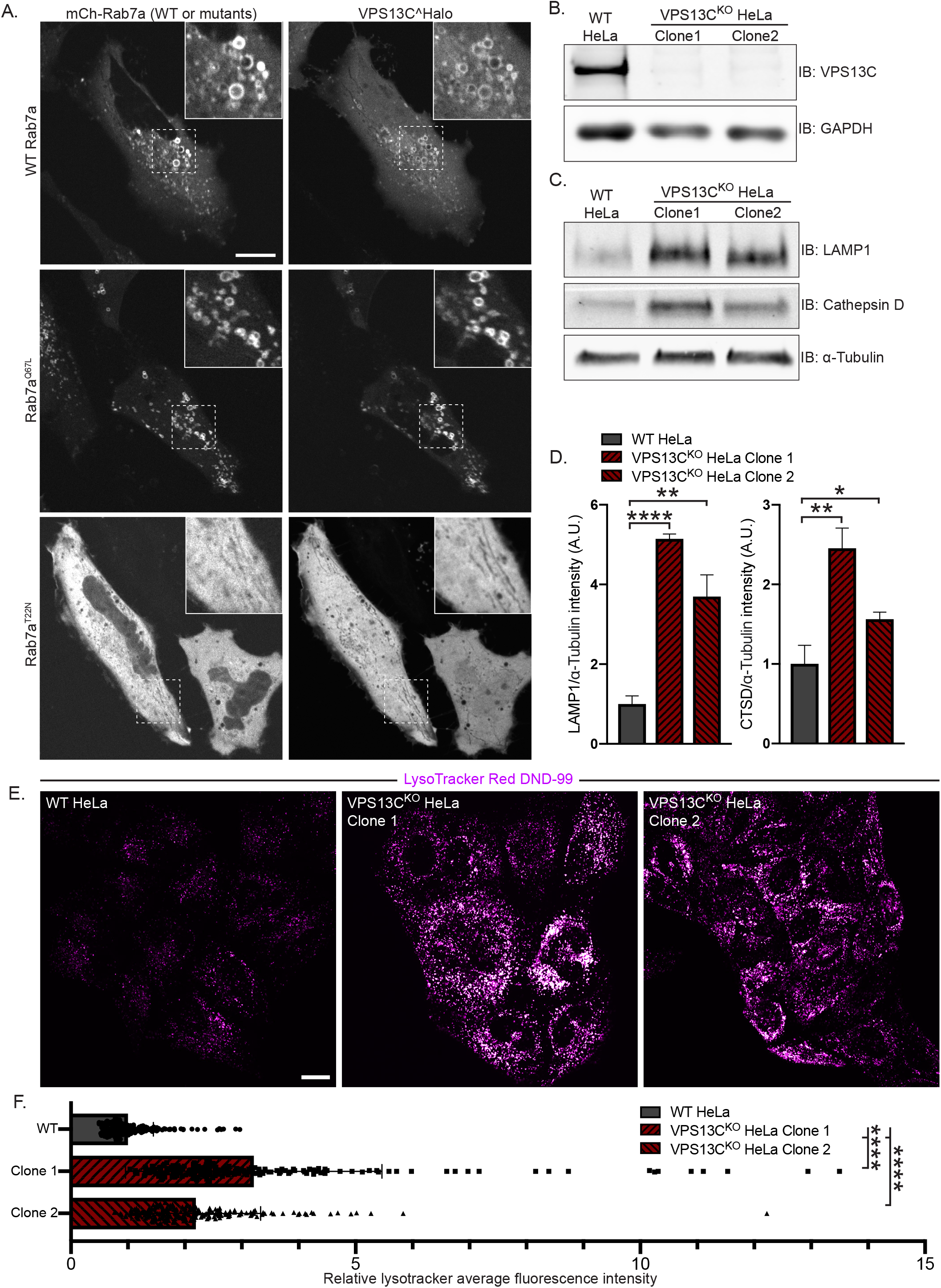
VPS13C is a Rab7 binding protein implicated in maintaining lysosomal homeostasis. (A) Live HeLa cells expressing full length VPS13C^Halo with either WT mCherry-Rab7a (top row), constitutively active mCherry-Rab7a^Q67L^ (middle row), or mCherry-Rab7a^T22N^ (bottom row). (B) Immunoblot of VPS13C in WT HeLa cells and two individual clonal cell lines after CRISPR-Cas9 mediated knockout of VPS13C. (C) Immunoblot of Lamp1 and Cathepsin D in WT and VPS13C^KO^ cells, quantified in (D). α-tubulin was used as a loading control. n=3 biological replicates. (E) Live WT and VPS13C^KO^ HeLa cells stained with LysoTracker Red DND-99 (50 nM), quantified in (F), n=3 biological replicates. Scale bars 20 μm. * P < 0.05, ** P < 0.01, **** P < 0.0001.

We generated two independent VPS13C^KO^ HeLa cell lines using CRISPR-Cas9 genome editing and confirmed indel mutations leading to premature stop codons by genomic sequencing (Figure S1A). Loss of protein expression was validated by immunoblotting (Figure 1B). Both VPS13C^KO^ clones had significantly elevated levels of the lysosomal membrane protein LAMP1 and of the luminal protease Cathepsin D, as assessed by western blotting (Figure 1C,D). Moreover, imaging assays revealed an increased lysotracker signal (Figure 1E,F), further supporting an increase in lysosome abundance and showing that these lysosomes have an acidic lumen.

Given the putative role of VPS13C as a lipid transfer protein, we next examined the impact of the absence of VPS13C on the lysosome lipidome. In order to isolate lysosomes, we pulsed (4 hrs) cells with dextran-coated superparamagnetic iron-oxide nanoparticles (SPIONs), which were taken up through bulk-endocytosis and trafficked to the endolysosomal compartment (Tharkeshwar et al., 2020; Tharkeshwar et al., 2017). Imaging of a fluorescently-tagged version of these nanoparticles at 15 hrs after the pulse confirmed their trafficking to vesicular structures which were positive for both LAMP1 and for a transfected construct comprising the beta-propeller region of VPS13C (VPS13C^b-prop^), i.e. the Rab7-binding region of VPS13C (Figure S2A). This observation confirmed the accumulation of the nanoparticles in a VPS13C-relevant compartment. Cell lysis and purification of nanoparticles-enriched lysosomes using a magnetic column yielded > 67-fold enrichment of the integral lysosomal membrane protein LAMP1 relative to the control protein GAPDH, as well as an enrichment of the late-endosomal marker Rab7, a peripheral membrane protein (Figure 2A,B).

**Figure 2.**
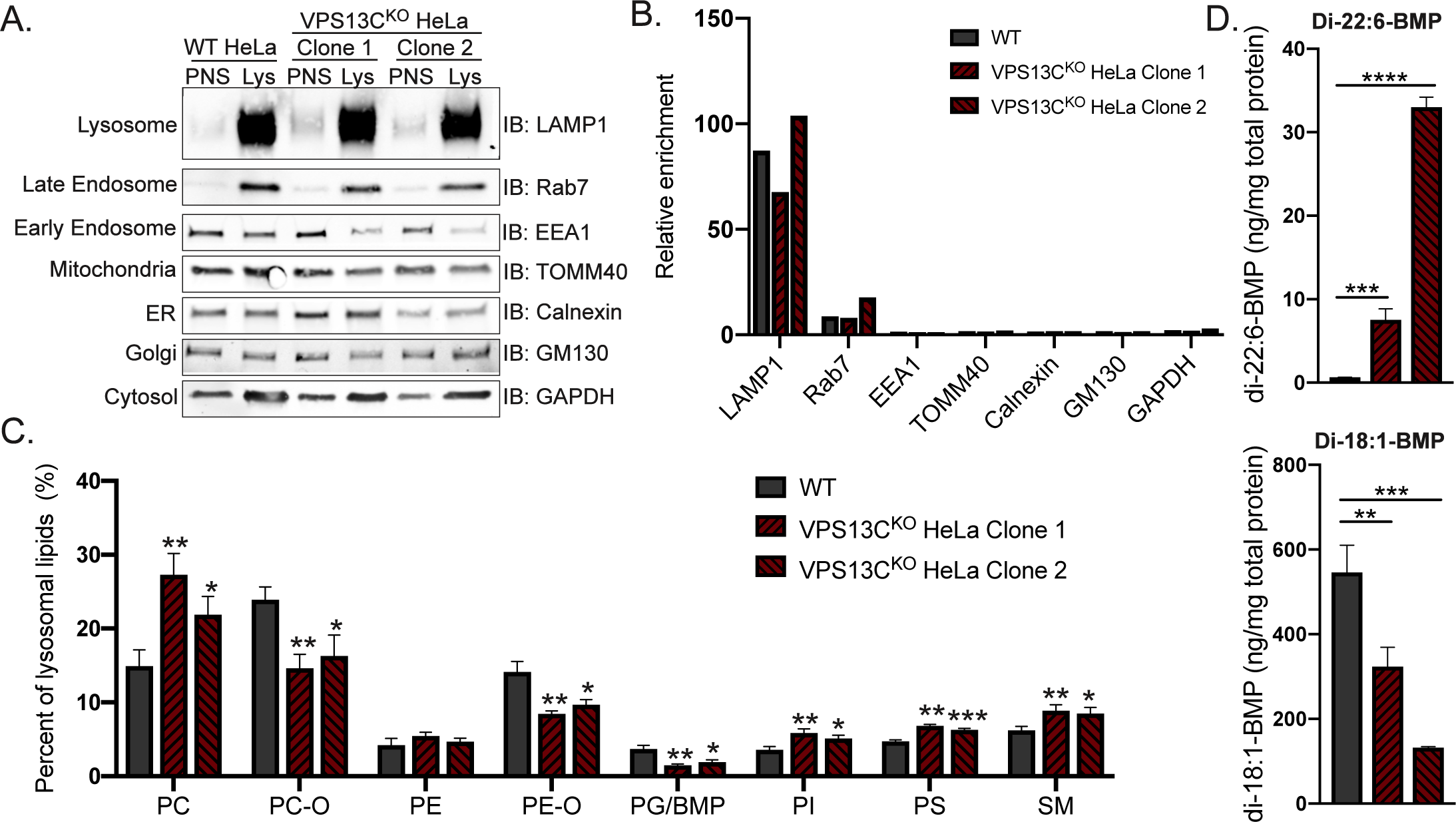
Loss of VPS13C results in altered lysosomal lipid composition. (A) Immunoblot showing abundance of organelle markers in post-nuclear supernatant (PNS) and lysosomal fractions (Lys) from WT and VPS13C^KO^ HeLa cells. Equal amounts of total protein was loaded in each lane. Note the striking enrichment of LAMP1 in lysosomal fractions. Quantification of band intensities from (A) normalized to GAPDH to show relative enrichment. (C) Percentages of phospho-and sphingolipid classes in WT and VPS13C^KO^ lysosomal fractions normalized to total measured lysosomal lipid content. N = 4 biological replicates. (D) Concentrations of di-22:6-and di-18:1-BMP normalized to total protein in WT and VPS13C^KO^ HeLa total cell lysate. N = 3 biological replicates. * P < 0.05, ** P < 0.01, *** P < 0.001, **** P < 0.0001 compared to WT control.

Shotgun mass spectrometry-based lipidomic analysis of the major lipid classes in the lysosomal fractions revealed substantial differences between VPS13C^KO^ and controls on a percent molar basis (specific lipid class versus total lipids) that were consistent in both VPS13C^KO^ clones. There were increases in phosphatidylcholine (PC), phosphatidylserine (PS), phosphatidylinositol (PI), and sphingomyelin (SM), as well as a decrease in phosphatidylglycerol (PG). PG measurement may include bis(monoacylglycerol)phosphate (BMP), as PG and BMP are structural isomers that were not distinguished by this analysis (Figure 2C). In addition, ether-lipid forms of both phosphatidylcholine (PC-O) and phosphatidylethanolamine (PE-O) were significantly reduced in the lysosomes of both VPS13C^KO^ clones (Figure 2C), though there was no decrease in these lipids, and even a slight increase, in the total cell lipidome (Figure S2B).

The whole cell lipidome also revealed increases in ceramide, hexosylceramide, and cholesteryl ester, as well as decreases in lysoPC (LPC) and lysoPE (LPE) (Figure S2B). Collectively, these findings reveal a perturbation of lysosomal lipid homeostasis in VPS13C^KO^ cells.

### Enhanced levels of di-22:6-BMP in VPS13C^KO^ cells

We next analyzed individual lipid species in control and VPS13C^KO^ lysosomes. Among the 1161 species measured, we found that 123 of them were significantly altered in both VPS13C^KO^ clones relative to controls (Figure S2C). In agreement with the class-level decreases, most of the downregulated hits were PC-O and PE-O species. The upregulated hits comprised a variety of classes including PC, PI, PE, and SM, many containing polyunsaturated fatty acid (PUFA) tails including arachidonic acid (20:4), eicosapentaenoic acid (20:5), docosapentaenoic acid (22:5), and docosahexaenoic acid (DHA, 22:6). The most highly increased lipid species in one of the VPS13C^KO^ clones and the third highest in the other one, was PG(22.6_22.6) (Figure S2C), black arrowhead), which, as stated above, could not be distinguished from its structural isomer di-22:6-BMP. As BMPs (also referred to as LBPA) are specific to the endolysosomal system (Gruenberg, 2020), we suspected that the majority of the species reported as PG(22.6_22.6) was actually di-22:6-BMP. This was intriguing, as di-22:6-BMP has been established as a biomarker for a number of neurodegenerative diseases, including Niemann Pick type C (Liu et al., 2014) and, more recently, LRRK2 G2019S mutation status (Alcalay et al., 2020). The increase of di-22:6-BMP in whole cell lysate was confirmed by a quantitative mass spectrometry assay specifically designed to assess this lipid (Figure 2D). Moreover, di-18:1-BMP, which appears to be the most abundant species in HeLa cells and is a major species in most human tissues(Showalter et al., 2020) was decreased (Figure 2D). This is consistent with our findings that total PG/BMP species are decreased by about half in VPS13C^KO^ lysosomes, revealing an overall decrease in total BMP but a specific increase in di-22:6-BMP (Figure 2C and S2D). Total BMP is also reduced in certain subtypes of Neuronal Ceroid Lipofuscinosis (NCL)(Hobert and Dawson, 2007), and is also altered by knockout of Progranulin, a lysosomal protein whose loss-of-function mutation cause both NCL (homozygous) and Frontotemporal Dementia (FTD) (heterozygous) (Logan et al., 2020). While the significance of the specific increase in di-22:6-BMP and decrease of total BMP remains unclear, this finding is consistent with alteration of lysosomal function across multiple neurodegenerative conditions.

### Activation of the cGAS/STING pathway

Having defined an impact of the lack of VPS13C on lysosomal homeostasis, we next addressed the hypothesis that the absence of VPS13C could result in an activation of the cGAS-STING pathway (Figure 3A). First, as a readout of potential cGAS-STING activation, we analyzed a subset of interferon-stimulated genes (ISGs) (IFIT1, IFIT3, ISG15, OASL and STAT1) by qPCR and saw increased expression in both VPS13C^KO^ HeLa clones (Figure 3B). This increase was no longer observed when cGAS (Figure 3F) or STING (Figure 3D) were knocked down by siRNA (Figure 3C). Knockdown of cGAS or STING also decreased ISG expression in the WT cells, suggesting some basal level of STING signaling in HeLa cells.

**Figure 3.**
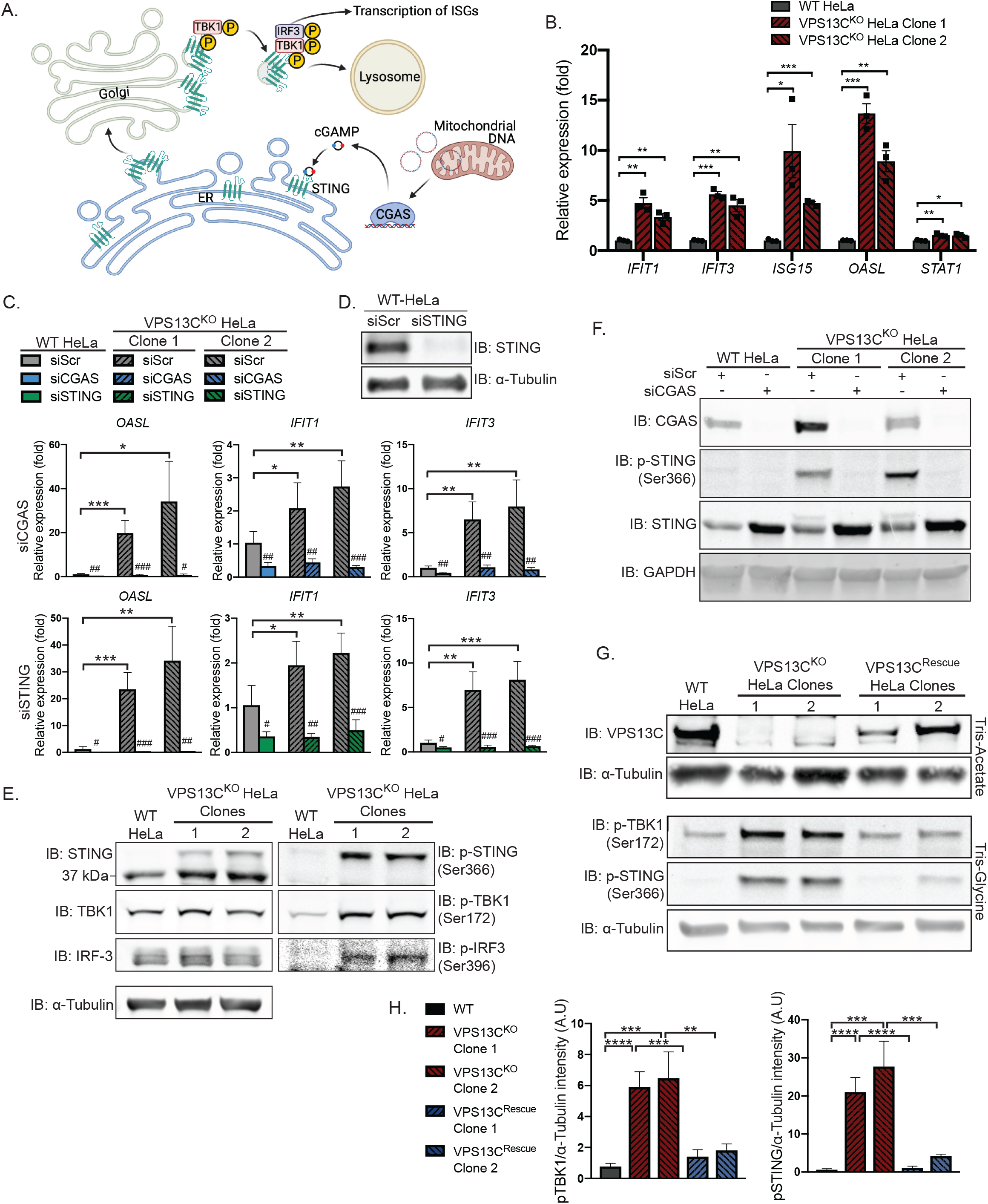
Loss of VPS13C results in activation of the cGAS/STING pathway. (A) Cartoon schematic of cGAS/STING signaling pathway and STING trafficking through the Golgi to lysosomes for degradation. Created with BioRender.com (B) qPCR of five ISG transcripts (*IFIT1, IFIT3, ISG15, OASL, STAT1*) shows increased expression in VPS13C^KO^ HeLa cells. N = 3 biological replicates. (C) qPCR of three ISG transcripts after treatment with siRNA against cGAS (top row) or STING (bottom row). N = 4 biological replicates. (D) Immunoblot showing efficiency of STING depletion after treatment with anti-STING siRNA. (E) Immunoblot showing increased levels of phosphorylated STING, TBK1, and IRF3, indicating activation of the cGAS/STING pathway. Note that the upper band in lanes 2 and 3 of the anti-STING blot corresponds to p-STING (lanes 2 and 3 of the p-STING blot. (F) Treatment of siRNA against cGAS significantly depletes cGAS and also returns p-STING to WT levels in the VPS13C^KO^ clones. cGAS-knockdown also causes an increase in total STING levels in both WT and VPS13C^KO^ cells. (G) Immunoblot showing lack of VPS13C band in VPS13C^KO^ cells and return of band in repaired VPS13C^Rescue^ clones. P-TBK1 and p-STING are returned toward WT levels in VPS13C^Rescue^ clones, quantified in (H). N = 3 biological replicates. * P < 0.05, ** P < 0.01, *** P < 0.001, **** P < 0.0001. # P < 0.05, ## P < 0.01, ### P < 0.001 in siCGAS or siSTING compared to siScr treated cells.

Upon binding to cGAMP, activated STING undergoes a conformational change, oligomerizes, traffics through the Golgi, and recruits the kinase TBK1, which phosphorylates STING at ser366 as well as itself at ser172(Liu et al., 2015; Shang et al., 2019; Zhang et al., 2019). Phosphorylated STING subsequently recruits IRF3, which is phosphorylated and activated by TBK1(Liu et al., 2015) and undergoes translocation to the nucleus to induce ISG expression (Figure 3A)(Lin et al., 1998). Again consistent with cGAS-STING activation, phosphorylated forms of STING, TBK1, and IRF3 were all significantly elevated in VPS13C^KO^ HeLa cells (Figure 3E). Total STING levels were also slightly increased (Figure 3E). Additionally, siRNA knockdown of cGAS abolished STING-Ser366 phosphorylation in VPS13C^KO^ HeLa cells, while global levels of STING were slightly increased consistent with lower basal degradation in the absence of cGAS (Figure 3F).

We next investigated whether activation of cGAS/STING could be rescued by VPS13C expression. Because transfected plasmid DNA can activate cGAS, we used CRISPR-Cas9 mediated homology directed repair (HDR) to correct one of the mutant VPS13C alleles in each of the VPS13C^KO^ clones (Figure 3G). In clones in which VPS13C protein expression had been successfully restored (VPS13C^Rescue^), phospho-STING and phospho-TBK1 were also restored to WT or near WT levels (Figure 3G,H).

### Role of mtDNA escape in cGAS/STING activation in VPS13C^KO^ cells

Since mtDNA is a ligand for cGAS and hence an activator STING (West et al., 2015), and mutations in the single yeast Vps13 protein results in escape of mtDNA(Thorsness and Fox, 1993), we considered the possible role of mtDNA leakage in the activation of STING observed in VPS13C^KO^ cells. While VPS13C localization does not suggest a direct impact of this protein on mitochondrial function, recent studies have revealed that alteration of lysosome function can indirectly impact mitochondria (Hughes et al., 2020; Kim et al., 2021; Yambire et al., 2019).

Moreover, mitochondrial defects were reported in Cos7 cells upon siRNA-mediated VPS13C knockdown (Lesage et al., 2016). To assess whether STING activation in VPS13C^KO^ cells was dependent on mtDNA, we treated VPS13C^KO^ cells with ethidium bromide (EtBr) for 72 hours to deplete mtDNA (Khozhukhar et al., 2018). The efficacy of this treatment was verified by qPCR of the D-loop region of mtDNA, which demonstrated a >97% depletion in mtDNA levels (Figure S3A). Depletion of mtDNA in VPS13C^KO^ cells reversed both the elevated expression of ISGs and the increased levels of phospho-STING and phospho-TBK1 (Figure 4A,B). Consistent with these results, we also observed a 2-4 fold increase in mtDNA (Figure 4C) in cytosolic fractions of VPS13C^KO^ cells, in which absence of mitochondria was documented by immunoblotting for the mitochondrial marker HSP60 (Figure S4B). However, no gross differences in either mitochondrial morphology or nucleoid structure or distribution in VPS13C^KO^ cells were observed by immunofluorescence (Figure S3C). Together these results support the hypothesis that cytosolic mtDNA is a major source of cGAS/STING activation in VPS13C^KO^ cells.

**Figure 4.**
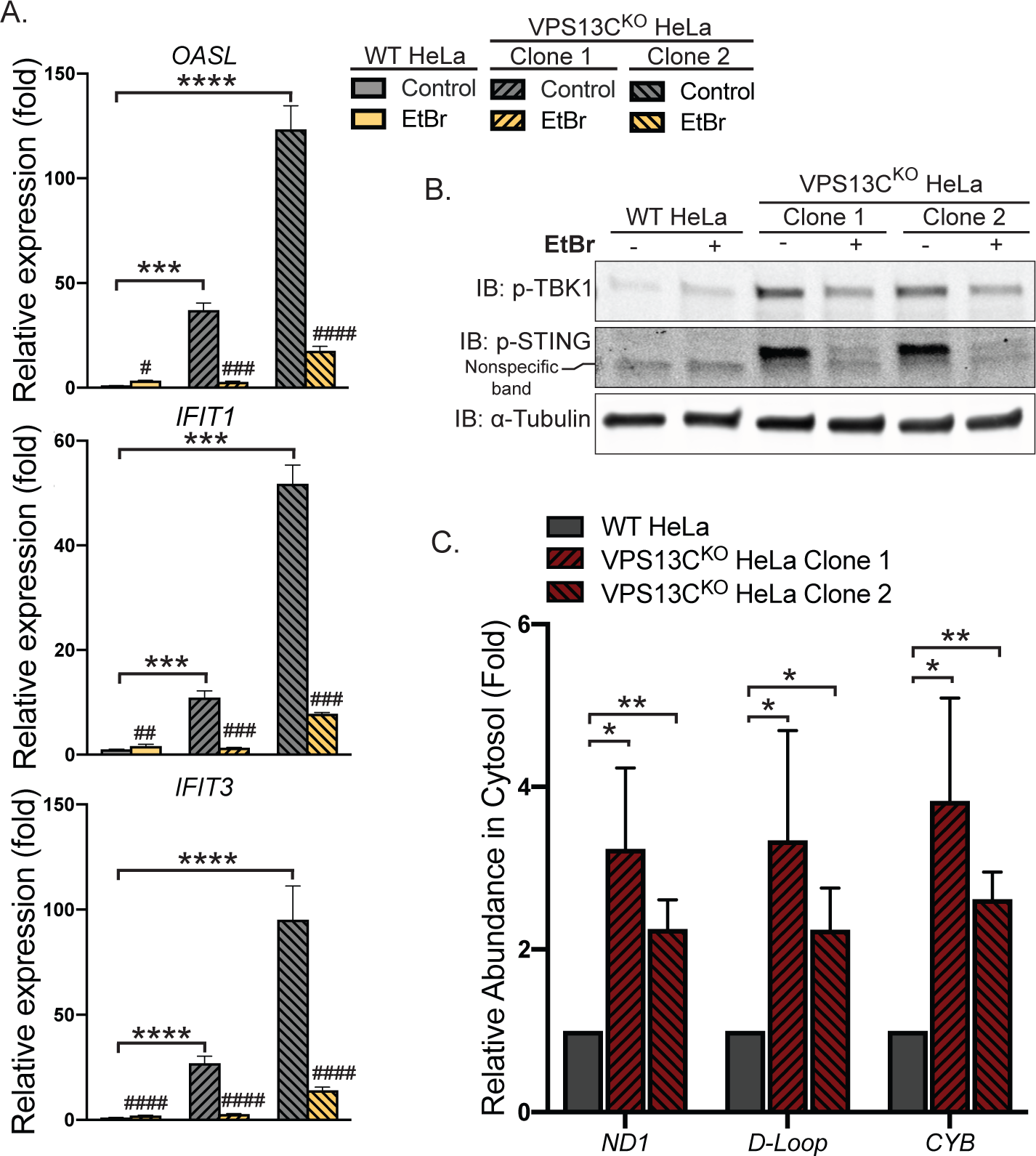
Activation of the cGAS/STING pathway in VPS13C^KO^ cells is dependent on increased cytosolic mtDNA. (A) qPCR of three ISG transcripts shows that depletion of mtDNA with EtBr reduces ISG levels in VPS13C^KO^ cells to or near WT levels. N = 3 biological replicates. (B) p-TBK1 and p-STING are reduced in EtBr treated VPS13C^KO^ cells. (C) Levels of three mtDNA amplicons (ND1, D-Loop, CYB) are elevated in the cytosolic fraction of VPS13C^KO^ cells. N = 3 biological replicates. * P < 0.05, ** P < 0.01, *** P < 0.001, **** P < 0.0001. # P < 0.05, ## P < 0.01, ### P < 0.001, #### < 0.0001 in EtBr compared to untreated cells.

### Steady-state change in the localization of STING in VPS13C^KO^ cells

The cGAMP-dependent oligomerization of STING leading to its activation also triggers its transport from the ER via the Golgi complex to lysosomes, where it is degraded leading to termination of signaling (Gonugunta et al., 2017; Gui et al., 2019). Thus, we tested whether the constitutive activation of STING observed in VPS13C^KO^ cells is accompanied by a change in its steady-state localization. As transient transfection, which involves acute introduction of plasmid DNA, can activate cGAS/STING, we generated cell lines stably expressing STING-GFP in control and VPS13C^KO^ HeLa cells via retroviral transduction. STING-GFP expression was similar in control and VPS13C^KO^ cells (Figure S4A), and phospho-STING(Ser366) and phospho-TBK1(Ser172) remained elevated in the VPS13C^KO^ cells (Figure S4B,C). In WT cells STING-GFP was almost exclusively localized to the ER, as expected (Figure 5A,B). Treatment with cGAMP, the product of cGAS that binds to STING, caused STING to concentrate in a Golgi complex-like pattern within 15 minutes and then to disperse throughout the cells as punctate structures, previously shown to overlap in part with lysosomes over the next six hours (Figure 5A, Video 1)(Gui et al., 2019). In contrast, in VPS13C^KO^ cells (not exposed to exogenous cGAMP), STING-GFP already had a predominantly punctate localization (Figure 5B,C) similar to the localization of STING-GFP in WT cells at 12 hrs after cGAMP stimulation (Figure 5D,E).

**Figure 5.**
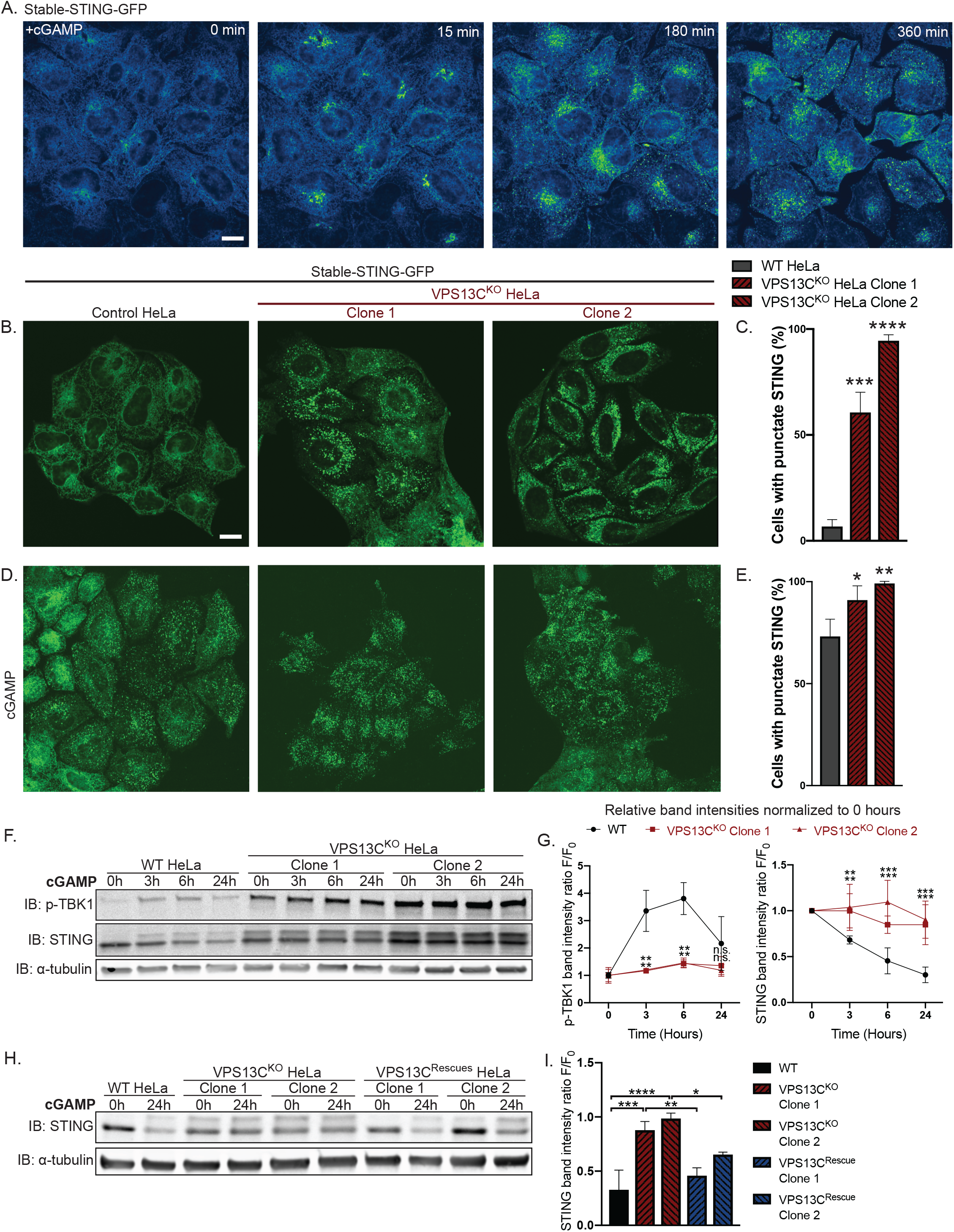
STING is activated and translocated out of the ER at baseline in VPS13C^KO^ cells. (A) Selected frames from timelapse of stable STING-GFP in WT HeLa cells after treatment with 50 ug/mL cGAMP (Video 1). STING-GFP is localized in an ER-like pattern at 0 min post-treatment, traffics to a Golgi-like pattern 15 min post-treatment, a Golgi/vesicular pattern by 180 minutes post-treatment, and a largely vesicular pattern by 360 min. (B) Under unstimulated basal conditions, STING-GFP is localized in an ER distribution in WT HeLa cells but a vesicular distribution in the majority of VPS13C^KO^ cells. The percentage of cells with vesicular pattern is quantified in (C). N = 3 biological replicates. (D) treatment with cGAMP had only minimal effect on the already punctate distribution of STING-GFP in VPS13C^KO^ cells, but induces a vesicular pattern in the majority of WT cells, quantified in (E). N = 3 biological replicates. (F) In WT cells, treatment with 8 ug/mL cGAMP causes an increase in p-TBK1 at 3h and 6h timepoints and a return to baseline at 24h, along with a concomitant decrease in total STING levels over 24 hours as STING is degraded. In VPS13C^KO^ cells, the same treatment fails to cause a significant increase in p-TBK1, STING upper band (phospho-STING), or decrease in total STING levels. (G) Band intensity of p-TBK1 and STING at each timepoint is quantified relative to the 0h value for each cell line. N = 3 biological replicates. (H) Treatment with 8 ug/mL cGAMP in VPS13C^Rescue^ cells for 24h results in STING degradation closer to WT levels, but fails to induce significant STING degradation in VPS13C^KO^ cells, quantified in (I). N = 3 biological replicates. Scale bars 20 μM. * P < 0.05, ** P < 0.01, *** P < 0.001, **** P < 0.0001 compared to WT control.

We complemented these localization studies with biochemical experiments. In WT cells, addition of herring testes (HT)-DNA to activate cGAS (Figure S4D), or of cGAMP to directly activate STING (Figure 5F) resulted in the appearance of the upper STING band (the phosphorylated form) and increased levels of phospho-TBK1, but also in the degradation of total STING over time, as reported (Gonugunta et al., 2017; Gui et al., 2019). After 24 hrs, total levels of STING were reduced to 25% (for HT-DNA)(Figure S4D,E) and 30% (for cGAMP)(Figure 5F,G) of baseline and phospho-TBK1 also returned towards baseline (Figure 5F,G). In contrast, in VPS13C^KO^ cells both phospho-TBK1 and phospho-STING were already elevated at baseline (total STING was also elevated), and addition of cGAMP did not promote degradation of these proteins (Figure 5F,G). These differences from WT were rescued in the VPS13C^Rescue^ cells where the VPS13C mutation had been repaired (Figure 5H,I). Collectively, these findings, i.e. elevated levels of both total STING and phospho-STING, and lack of its responsiveness to stimulation, raised the questions of whether the cGAS-STING pathway could still be activated by exogenous cGAMP in VPS13C^KO^ cells and whether degradation of STING might also be impaired.

### Silencing of cGAS unmasks cGAMP responsiveness of VPS13C^KO^ cells and reveals impaired STING degradation

To determine whether VPS13C^KO^ cells can still respond to cGAMP, these cells were treated with siRNA against cGAS (with scrambled siRNA as a control), to suppress basal STING activation. cGAS knock-down reverted STING-GFP to an ER localization (Figure 6A,B), similar to untreated WT cells, while scrambled siRNA had no effect (Figure 6A,C).

**Figure 6.**
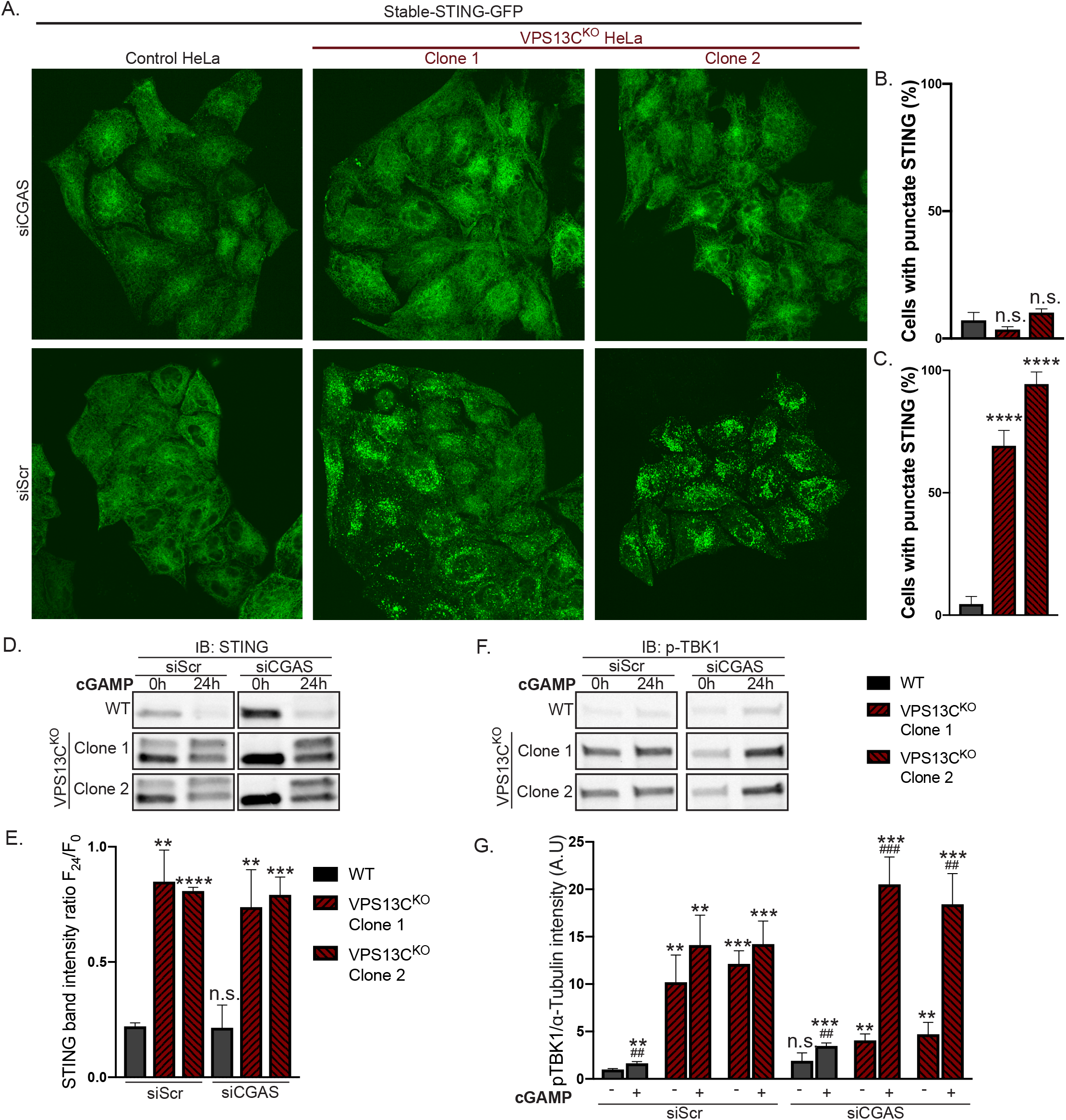
Silencing of cGAS unmasks cGAMP responsiveness and reveals impaired STING degradation in VPS13C^KO^ cells. (A) Treatment with siRNA against cGAS returns STING-GFP to an ER distribution in VPS13C^KO^ cells, similar to untreated WT cells, quantified in (B), while siScr has no effect, quantified in (C). N = 3 biological replicates. (D) Immunoblot against STING in WT and VPS13C^KO^ cells treated with either scrambled siRNA or siRNA against CGAS, followed by treatment with 8 ug/mL cGAMP for 24h. Treatment with siRNA against CGAS returns STING to the unphosphorylated state in VPS13C^KO^ cells, rendering them responsive to cGAMP. In VPS13C^KO^ cells, the activation of STING is not followed by degradation, compared to WT cells. (E) Quantification of the total STING signal remaining at 24h after cGAMP treatment (F24) relative to 0h (F0). (F) Immunoblot against p-TBK1 under the same conditions as (E). WT cells behaved similarly in response to cGAMP treatment regardless of siScr or siCGAS pretreatment, with p-TBK1 levels returning to baseline after 24h of cGAMP treatment, as shown by the timecourse of Figure 5F and G. In VPS13C^KO^ cells pretreated with siScr, p-TBK1 remained at baseline after 24h of cGAMP treatment, presumably never having increased, based on the timecourse in Figure 5F and G. In VPS13C^KO^ cells pretreated with siCGAS however, p-TBK1 was significantly elevated after 24h of cGAMP, in accordance with the defect in STING degradation and continued STING signaling (E and F). (G) Quantification of p-TBK1 band intensity. N = 3 biological replicates. ** P < 0.01, *** P < 0.001, **** P < 0.0001 compared to untreated WT cells. ## P < 0.01, ### P < 0.001 value at 24h cGAMP treatment compared to corresponding 0h cGAMP treatment.

Biochemical experiments confirmed the restoration of responsiveness to cGAMP of VPS13C^KO^ cells. Both phospho-TBK1 (Figure 6F,G), and phospho-STING (Figure S5B), (also reflected by the upper total STING band (Figure 6D)) returned to unstimulated WT levels following treatment with anti-cGAS siRNA (Full blot in Figure S5A). Upon addition of cGAMP, we observed only a very modest reduction of total STING levels at 24h in VPS13C^KO^ cells compared to significant STING degradation in WT cells, suggesting a bona fide defect in STING degradation (Figure 6D,E). Furthermore, phospho-TBK1(Figure 6F,G) and phospho-STING (Figure S5B) remained elevated in VPS13C^KO^ cells at 24h, in agreement with sustained STING signaling.

The defect in STING degradation did not reflect an overall defect in protein degradation in lysosomes, as we found no difference in the kinetics of EGFR degradation in response to EGF stimulation between VPS13C^KO^ and WT cells (Figure S6A,B). Likewise, we found no difference in LC3 degradation after induction of macro autophagy by starving cells in EBSS (Figure S6C). Interestingly, the ratio of lipidated LC3-II to LC3-I under basal conditions was increased in the VPS13C^KO^ cells (Figure S6C), consistent with the reported property of activated STING to induce LC3 lipidation(Fischer et al., 2020; Gui et al., 2019).

## DISCUSSION

Our results support a role of VPS13C in regulating lysosome function and show that cellular perturbations produced by the absence of VPS13C result in activation of the cGAS-STING pathway. Such activation is of special interest as VPS13C is a PD gene and aberrant activation the cGAS-STING pathway has been implicated in PD pathogenesis(Sliter et al., 2018).

We have previously shown a colocalization of VPS13C at the interface between the ER and organelles positive for Rab7, a marker of late endosomes and lysosomes. Moreover, VPS13C was identified in a screen for Rab7 effectors(Gillingham et al., 2019). Our present finding that dominant negative Rab7 completely blocks VPS13C recruitment to lysosomes proves its major role in controlling VPS13C localization. Importantly, we show that absence of VPS13C results in alterations in lysosome homeostasis, as demonstrated by an increase in the levels of lysosome markers and by alterations of the lipid composition of purified lysosomal fractions. Increases were observed in PC, PI, PS, and SM, with decreases in ether-linked phospholipids (PC-O and PE-O) and in BMP, a class of lipids characteristic of multivesicular bodies and lysosomes (Gruenberg, 2020). Whether these changes are directly or indirectly related to the property of VPS13C to transport lipids remains to be explored, as do the functional implications of these changes. Of note, however, was the robust and highly specific accumulation of di-22:6-BMP in both lysosomes and total cell lysates, in spite of the overall reduction of BMP, as a specific increase in urinary di-22:6-BMP was reported to be a marker of LRRK2 G2019S mutation status (Alcalay et al., 2020). Like the absence of VPS13C, the LRRK2 G2019S mutation increases PD risk (West et al., 2005). It will be interesting to determine whether elevated di-22:6-BMP is causally linked to PD pathogenesis or is simply a marker of more general lysosome dysfunction.

Our finding that the cGAS/STING signaling pathway is activated in VPS13C^KO^ cells adds to evidence for a potential involvement of this pathway in PD pathogenesis, as first suggested by studies of PINK1 and Parkin mouse models (Sliter et al., 2018). The role of activation of this pathway in PD, and more generally in neurodegenerative diseases, is an area of intense investigation (McCauley et al., 2020; Sliter et al., 2018; Weindel et al., 2020; Yu et al., 2020). Our results suggest that the increase of STING signaling may result both from a leakage of mtDNA, which in turn would activate cGAS and thus generation of the STING ligand cGAMP, and from a delayed degradation of activated STING in VPS13C^KO^ cells. The mechanisms underlying this delayed degradation remain elusive. Presumably, STING must remain facing the cytosol (and not be internalized in the lysosomal lumen) to continue activating TBK1 and IRF3. A defective or incomplete fusion of STING-positive vesicles with late endosomes/lysosomes or a defect in the incorporation of STING into intraluminal vesicles of late endosomes are potential mechanisms. We speculate that alterations to the lysosomal lipidome may be responsible for these defects. Interestingly, defective lysosomal degradation of activated STING, leading to higher levels of innate immune signaling, was reported in cells deficient in C9orf72, another neurodegeneration gene associated with lysosomes (McCauley et al., 2020)

An attractive unifying scenario is that alteration of lysosome function is the most proximal consequence of VPS13C depletion, and that such alteration is upstream of mitochondrial dysfunction and STING activation. More specifically, defective lysosomal function may be the primary event leading to mtDNA leakage. A similar scenario may apply to STING activation by the absence of LRRK2, another PD protein implicated in lysosome function (Bonet-Ponce et al., 2020; Weindel et al., 2020). Indeed, it is now appreciated that genetic or pharmacologic disruption of lysosome function can lead to mitochondria dysfunction in a number of contexts (Hughes et al., 2020; Kim et al., 2021; Yambire et al., 2019). This cross-talk may be mediated by soluble factors (Hughes et al., 2020; Yambire et al., 2019) or by direct mitochondria lysosome contacts (Wong et al., 2018). It is also possible that leakage of mtDNA may occur during mitophagy by defective lysosomes. While mtDNA can escape directly from mitochondria into the cytosol (Riley and Tait, 2020; West and Shadel, 2017), as in the case of TFAM deficiency(West et al., 2015) or TDP-43 mutations (Yu et al., 2020), it may also escape during the process of mitophagy/autophagy (Gkirtzimanaki et al., 2018; Oka et al., 2012).

An important question for future studies will be the elucidation of how these results relate to VPS13C loss-of-function in brain cells. VPS13C is ubiquitously expressed in cells of the central nervous system. In contrast, cGAS and STING are primarily expressed in microglia (Saunders et al., 2018). Interestingly, several other genes relevant to neurodegeneration are primarily expressed in microglia (Hickman et al., 2018). We have attempted to differentiate human iPS VPS13C^KO^ cells into microglia to assess potential defects resulting from the absence of VPS13C. However, while the differentiation protocol used was successful in yielding WT microglial cells, it failed to yield VPS13C^KO^ microglial cells, revealing a possible importance of VPS13C in microglia. So far we have not observed obvious defects in VPS13C^KO^ mice, a finding that also applies to other PD genes, including PINK1 and Parkin (Sliter et al., 2018).

In conclusion, we have shown that in a model human cell line the absence of VPS13C results in late-endosome/lysosomal defects, as had been predicted by the localization of VPS13C at contacts between the ER and lysosomes and by the proposed role of VPS13C in mediating lipid exchange between these two organelles(Kumar et al., 2018; Leonzino et al., 2021). We have further discovered that these defects correlate with abnormally elevated STING signaling, most likely due to direct and indirect effects of the perturbation of lysosome function. While it remains to be seen how these findings can be generalized to other cell types and to intact tissues, our findings provide further support for the hypothesis that activation of innate immunity may be one of the factors involved in pathogenetic mechanisms resulting from mutations in PD genes.

## METHODS

### DNA plasmids

A plasmid containing codon-optimized cDNA encoding human VPS13C, with a Halo protein flanked by SacII restriction enzyme sites after amino acid residue 1914, was generated by and purchased from GenScript Biotech. mCherry-Rab7a was obtained from Addgene (#68104). mCherry-Rab7a^Q67L^ and mCherry Rab7a^T22N^ were generated in our lab as previously reported (Guillen-Samander et al., 2019). Lamp1-mGFP was obtained from Addgene (#34831). GFP-VPS13C^bprop^ was generated in our lab as previously described (Kumar et al., 2018). For CRISPR mediated gene editing, candidate guide RNAs (gRNAs) against the human VPS13C genomic locus were identified using Benchling. gRNAs were ordered as complementary single-stranded oligonucleotides from Integrated DNA Technologies (IDT) then cloned into the px459 plasmid (Addgene plasmid #62988) using a one-step ligation protocol (Ran et al., 2013), and gRNAs were sequence verified using the U6 forward promoter. For CRISPR repair of the mutated VPS13C locus, gRNA was ordered (IDT) that incorporated a single nucleotide insertion present in one VPS13C allele in both of the VPS13C^KO^ clones. This was again cloned into the px459 plasmid (Addgene plasmid #62988) using a one-step ligation protocol (Ran et al., 2013). To generate pMX-STING-GFP, STING-V1 plasmid was obtained from Addgene (Addgene #124262) and the STING coding sequence was amplified by PCR and ligated into pEGFP-N1 by using XhoI and SacII restriction sites. The STING-GFP coding sequence was then cut from the pEGFP-N1 backbone and ligated into a pMXs-IRES-Blasticidin Retroviral Vector backbone (RTV-016, Cell Biolabs) using XhoI and SacII restriction sites. Oligos used in this study are listed in Table S1. All oligos were purchased from IDT.

### Antibodies

Primary antibodies used: mouse α-tubulin (T5168, Sigma-Aldrich), rabbit calnexin (ADI-SPA-860-D, Enzo), rabbit cathepsin D (Ab75852, Abcam), rabbit CGAS (D1D3G, CST), mouse DNA (CBL186, EMD Millipore), rabbit EEA1 (PA1063A, Thermo), mouse GAPDH (1D4, Enzo), mouse GM130 (610822, BD), rabbit HSP60 (12165S, CST), rabbit IRF3 (D83B9, CST), mouse LAMP1 (H4A3, DSHB), rabbit Phospho-IRF3 (S396) (4D4G, CST), rabbit Phospho-STING (S366) (D7C3S, CST), rabbit Phospho-TBK1 (S172) (D52C2, CST), rabbit Rab7 (D95F2, CST), rabbit STING (D2P2F, CST), rabbit TBK1 (D1B4, CST), rabbit VPS13C (custom, Proteintech)

### Cell culture and transfection

HeLa-M cells were cultured at 37°C in 5% CO2 and DMEM containing 10% FBS, 100 U/ml penicillin, 100 mg/ml streptomycin, and 2 mM L-glutamine (all from Gibco). For live-cell imaging experiments, cells were seeded on glass-bottomed dishes (MatTek) at a concentration of 35,000 cells per dish and transfected after 24h using FuGene HD (Promega). For biochemical experiments, cells were plated at such density so as to be approximately 90% confluent at the time of lysis. Transfection of siRNA was accomplished using Lipofectamine RNAiMax (ThermoFisher). Introduction of cGAMP was accomplished using Lipofectamine RNAiMax (ThermoFisher) as previously reported (Swanson et al., 2017). Transfection of HT-DNA was accomplished using Lipofectamine 2000 (ThermoFisher). Plat-A cells for retroviral packaging were cultured at 37°C in 5% CO2 and DMEM containing 10% FBS, 1 100 U/ml penicillin, 100 mg/ml streptomycin, 2 mM L-glutamine, 1 μg/mL puromycin, and 10 μg/mL blasticidin.

### Generation of Stable STING-GFP cells using retrovirus

For retroviral packaging, 5 × 10^6^ Plat-A cells (Cell Biolabs) were plated on a 10 cm plate in media without antibiotics. The following day, cells were transfected with 9 µg of pMX-STING-GFP using Fugene HD (Promega). Retroviral supernatant was collected 72 hours post transfection, supplemented with Polybrene (8 μg/mL), and passed through 0.22 μm filter to remove cellular debris before being added to WT and VPS13C^KO^ HeLa cells. After 24 hours, retroviral supernatant was removed and replaced with fresh DMEM complete. After an additional 24 hours, HeLa cells were FACS sorted to enrich for GFP positive cells. pMX-STING-GFP

### Generation of CRISPR-KO and CRISPR-KI rescue cell lines

Early passage HeLa-M cells were transfected with 1.5 µg of px459 plasmid (Addgene plasmid #62988) containing a single guide RNA (sgRNA) against VPS13C using Lipofectamine 2000 (ThermoFisher). At 24h post-transfection, cells were selected in complete DMEM containing 2 μg/mL puromycin. At 48h and 72h post transfection, media was replaced with fresh puromycin-containing media. After three days of puromycin selection, single clones were obtained using serial dilution and then screened by immunoblot. Two clonal cell lines lacking VPS13C by immunoblot were selected. Genomic DNA was extracted using a DNAeasy kit (Qiagen) and a ∼500 bp portion around the predicted CRISPR cut site was amplified by PCR and purified by NucleoSpin Gel and PCR Clean-up kit (Macherey-Nagel). This fragment was then ligated into an EGFP-N vector using Xho1 and Apa1 restriction sites and transformed into DH5-a competent cells. After plating the transformation mix onto agar plates, >48 bacterial colonies per clone were submitted for Sanger sequencing to maximize the probability of sequencing all alleles. Frameshift mutations leading to premature stop codons were identified by Sanger sequencing.

To rescue VPS13C expression we used CRISPR mediated homology directed repair (HDR) using a single-stranded oligo DNA nucleotide (ssODN) as previously described (Okamoto et al., 2019). Briefly, a gRNA was synthesized that incorporated a single nucleotide insertion present in one VPS13C allele in both of the VPS13C^KO^ clones. An ssODN corresponding to the WT VPS13C sequence flanking the insertion was generated. Both VPS13C^KO^ clones were transfected with 1μg of px459 plasmid containing the gRNA against the mutant VPS13C allele and 100 pmols of ssODN. Puromycin selection and single clone selection was performed as above, and rescue of protein expression was confirmed by immunoblot.

### Light microscopy

#### Live-cell imaging

Prior to imaging, growth medium was removed and replaced with live-cell imaging solution (Life Technologies). All live-cell imaging was performed at 37°C in 5% CO2. Imaging was performed using an Andor Dragonfly spinning-disk confocal microscope equipped with a plan apochromat objective (63x, 1.4 NA, oil) and a Zyla scientific CMOS camera. For lysotracker experiments, cells were incubated in 50 nM LysoTracker Red DND-99 (ThermoFisher) in complete DMEM for 30 minutes, washed twice with media, then imaged in live-cell imaging solution.

#### Immunofluorescence

WT and VPS13C^KO^ HeLa cells were plated on glass coverslips and fixed in a pre-warmed (37°C) solution of 4% paraformaldehyde in PBS for 15 minutes at room temperature, permeabilized with 0.1% (v/v) Triton X-100 in PBS for 10 minutes at room temperature, and blocked using filtered PBS containing 1% (w/v) BSA for an hour at room temperature. Coverslips were then incubated with antibodies against DNA (CBL186, EMD Millipore, 1:150) and HSP60 (12165S, CST, 1:1000), diluted in filtered PBS containing 1% BSA at 4°C overnight, followed by 3 × 5-minute washes in PBS. Secondary antibodies (1:1000, Alexa fluorophores 488 and 555, Invitrogen) were incubated in PBS containing 1% BSA for 1 hour at room temperature and removed by 3 × 5-minute washes in PBS. Finally, coverslips were mounted onto slides using ProLong Gold Antifade Mountant with DAPI (ThermoFisher P36935) and allowed to cure overnight at room temperature prior to imaging.

#### Image processing and analysis

Fluorescence images were processed using Fiji software (ImageJ; National Institutes of Health). For quantification of lysotracker images (Figure 1E), cells were outlined manually and their average fluorescence intensity measured. Intensities were then normalized such that the average intensity of the WT cells was 1. For quantification of stable STING-GFP cells, cells with a reticular ER or vesicular pattern were counted manually and the percentage of cells with punctate STING-GFP distribution was calculated.

### Synthesis of colloidal dextran-conjugated superparamagnetic iron nanoparticles

**(SPIONs)** SPIONs were synthesized according to the following protocol: 10 mL of aqueous 1.2 M FeCl2 and 10 mL of 1.8 M FeCl3 were combined in a 500 mL glass beaker with magnetic stirring, followed by slow addition of 10 mL of 30% NH4OH. This mixture was stirred for 5 minutes and during this time a dark brown sludge formed. The beaker was then placed on a strong magnet to allow the particles to migrate towards the magnet. The supernatant was removed and the particles resuspended in 100 ml water, followed by repeated separation on the magnet. This was step was repeated two more times. Particles were then resuspended in 80 mL of 0.3 M HCl and stirred for 30 minutes, followed by addition of 4 g of dextran and another 30 minutes of stirring. Particles were transferred into dialysis bags and dialyzed for 48 hours in milliQ water with water changes approximately every 12 hours. The resulting mixture was then centrifuged at 19,000 g for 15 minutes to remove large aggregates. Supernatant containing nanoparticles was stored at 4°C.

### Purification of lysosomes with dextran-conjugated SPIONs

WT and VPS13C^KO^ HeLa cells were plated on 4x 15 cm plates at 3.5 × 10^6^ cells per dish. The following day, the culture medium (DMEM) was exchanged for fresh DMEM containing 10 mM HEPES and 10% SPION solution by volume for 4 hours (pulse). The medium was then changed back to fresh DMEM and after 15 hours (chase) the cells were washed twice with PBS and then scraped into 5 mL of PBS on ice. Cells were then centrifuged at 1000 rpm for 10 min at 4°C. PBS was removed and the cell pellet was resuspended in 3mL homogenization buffer (HB)(5mM Tris, 250 mM Sucrose, 1mM EGTA pH 7.4) supplemented with protease inhibitor cocktail (Roche) immediately before use) and passed through a manual cell homogenizer (Isobiotec, 10 cycles, 10-micron clearance) to generate a total cell lysate. The lysate was centrifuged at 800 g for 10 min at 4°C, and the supernatant was loaded onto a magnetic LS column (Miltenyi Biotec) pre-equilibrated with 1mL of HB. The column was washed with 5 mL HB, then removed from the magnetic rack and eluted with 3 successive aliquots of 1mL HB forced through with positive pressure. The eluate was then centrifuged at 55,000 rpm for 1 hr at 4°C to pellet the lysosome fraction and the resulting pellet was resuspended in 200 µL of mass-spectrometry grade water (ThermoFisher) and flash frozen. Fluorescence images of nanoparticles were obtained using FluoreMAG A ferrofluid (Liquids Research Ltd) with a 4 hour pulse and 15 hour chase.

### Lipid extraction for mass spectrometry lipidomics

Mass spectrometry-based lipid analysis was performed by Lipotype GmbH (Dresden, Germany) as described (Sampaio et al. 2011). Lipids were extracted using a two-step chloroform/methanol procedure (Ejsing et al. 2009). Samples were spiked with an internal lipid standard mixture containing: cardiolipin 16:1/15:0/15:0/15:0 (CL), ceramide 18:1;2/17:0 (Cer), diacylglycerol 17:0/17:0 (DAG), hexosylceramide 18:1;2/12:0 (HexCer), lyso-phosphatidate 17:0 (LPA), lyso-phosphatidylcholine 12:0 (LPC), lyso-phosphatidylethanolamine 17:1 (LPE), lyso-phosphatidylglycerol 17:1 (LPG), lyso-phosphatidylinositol 17:1 (LPI), lyso-phosphatidylserine 17:1 (LPS), phosphatidate 17:0/17:0 (PA), phosphatidylcholine 17:0/17:0 (PC), phosphatidylethanolamine 17:0/17:0 (PE), phosphatidylglycerol 17:0/17:0 (PG), phosphatidylinositol 16:0/16:0 (PI), phosphatidylserine 17:0/17:0 (PS), cholesterol ester 20:0 (CE), sphingomyelin 18:1;2/12:0;0 (SM), triacylglycerol 17:0/17:0/17:0 (TAG). After extraction, the organic phase was transferred to an infusion plate and dried in a speed vacuum concentrator. The 1st step dry extract was re-suspended in 7.5 mM ammonium acetate in chloroform/methanol/propanol (1:2:4, V:V:V) and the 2nd step dry extract in a 33% ethanol solution of methylamine in chloroform/methanol (0.003:5:1; V:V:V). All liquid handling steps were performed using Hamilton Robotics STARlet robotic platform with the Anti Droplet Control feature for organic solvents pipetting.

### Mass spectrometry data acquisition

Samples were analyzed by direct infusion on a QExactive mass spectrometer (Thermo Scientific) equipped with a TriVersa NanoMate ion source (Advion Biosciences). Samples were analyzed in both positive and negative ion modes with a resolution of Rm/z=200=280000 for MS and Rm/z=200=17500 for MSMS experiments, in a single acquisition. MSMS was triggered by an inclusion list encompassing corresponding MS mass ranges scanned in 1 Da increments (Surma et al. 2015). Both MS and MSMS data were combined to monitor CE, DAG and TAG ions as ammonium adducts; PC, PC O-, as acetate adducts; and CL, PA, PE, PE O-, PG, PI and PS as deprotonated anions. MS only was used to monitor LPA, LPE, LPE O-, LPI and LPS as deprotonated anions; Cer, HexCer, SM, LPC and LPC O-as acetate adducts.

### Data analysis and post-processing

Data were analyzed with in-house developed lipid identification software based on LipidXplorer (Herzog et al. 2011; Herzog et al. 2012). Data post-processing and normalization were performed using an in-house developed data management system. Only lipid identifications with a signal-to-noise ratio >5, and a signal intensity 5-fold higher than in corresponding blank samples were considered for further data analysis.

### Measurement of di-22:6-BMP and di-18:1-BMP

Targeted high resolution UPLC-MS/MS was used to accurately quantitate total di-22:6-BMP (the sum of its 3 isoforms). Total di-18:1-BMP was measured as well. Quantitation was performed by Nextcea, Inc. (Woburn, MA) using authentic di-22:6-BMP and di-18:1-BMP reference standards. Di-14:0-BMP was employed as an internal standard.

### Quantitative PCR

RNA was extracted using the RNeasy plus micro RNA extraction kit followed by reverse transcription using iScript cDNA synthesis Kit (Bio-Rad). Equal amounts of DNA and corresponding primers (Supplementary Table 1) were used for quantitative PCR (qPCR) using SYBR Green Master Mix. For each biological sample, two technical replicates were performed. Mean values were normalized against the β-actin threshhold cycle (Ct) value to calculate ΔCt. The ΔCt of each sample was then compared to the ΔCt of the WT sample to generate the ΔΔCt value. Relative expression was then analysed using the 2−ΔΔCt method and the relative fold change was plotted with the WT samples given a value of 1. To quantify mtDNA, DNA was extracted from total cell lysate and cytosolic fractions using DNeasy kit (Qiagen). DNA samples were each diluted 1:10 and corresponding primers (Supplementary Table 1) were used for quantitative PCR (qPCR) using SYBR Green Master Mix. Two technical replicates were performed for each biological sample and mean Ct values of mtDNA amplicons form cytosolic fractions were normalized against the corresponding total cell lysate hB2M (nuclear DNA control) Ct value. Relative copy number was determined by the 2−ΔΔCt method and the WT mtDNA abundance was given a value of 1.

### Statistical analyses

GraphPad Prism 8 software was used for statistical comparison of live cell imaging, immunoblot densitometry, qPCR, and BMP measurement. Student’s t-test was used to assess significant differences between groups. Statistical analysis for lipidomic data was performed using RStudio. Groups were compared using student’s t-test followed by adjustment for multiple comparisons using Benjamini–Hochberg methodology to control false discovery rate (FDR). Q-values were determined using the R package ‘‘qvalue,’’ and a significance threshold of 0.05 was used. qPCR

## AUTHOR CONTRIBUTIONS

WHC, SF, GS, and PDC contributed to conceptualization and funding acquisition. WHC and ZW performed investigation, data curation, and formal analysis. AT contributed to methodology and resources. WHC and PDC contributed to visualization and writing – original draft. All authors contributed to validation of data and writing – review & editing.

## ACKNOWLEDGMENTS

We thank M. Leonzino, J.H. Park, A. Guillen-Samander, A. Iwasaki, R. Medzhitov and J. Gruenberg for discussion, A. Guillen-Samander for critical reagents, F. Wilson and A. Dao for outstanding technical assistance, and F. Hsieh and Nextcea Inc. for mass-spectrometry measurement of BMP species.

This work was supported in part by NIH grants NS36251 and DA018343 to P. De Camilli, the Parkinson Foundation (PF-RCE-1946) to P. De Camilli and S. Ferguson, NIH GM105718 to S. Ferguson, and NIH R01 AR069876 to G. Shadel. W. Hancock-Cerutti was supported in part by by NIH Medical Scientist Training Program Training Grant T32GM007205 and by NIH F31NS110229-01. Z.W. was supported by the China Scholarship Counsel. The study was also funded by the joint efforts of the Michael J. Fox Foundation for Parkinson’s Research (MJFF) and the Aligning Science Across Parkinson’s (ASAP) initiative. MJFF administers the grant ASAP-000580 on behalf of ASAP and itself. For the purpose of open access, the author has applied a CC-BY public copyright license to the Author Accepted Manuscript (AAM) version arising from this submission.

P. De Camilli serves on the scientific advisory board of Casma Therapeutics. The authors declare no further potential competing financial interests.

## FIGURE LEGENDS

**Figure S1.**
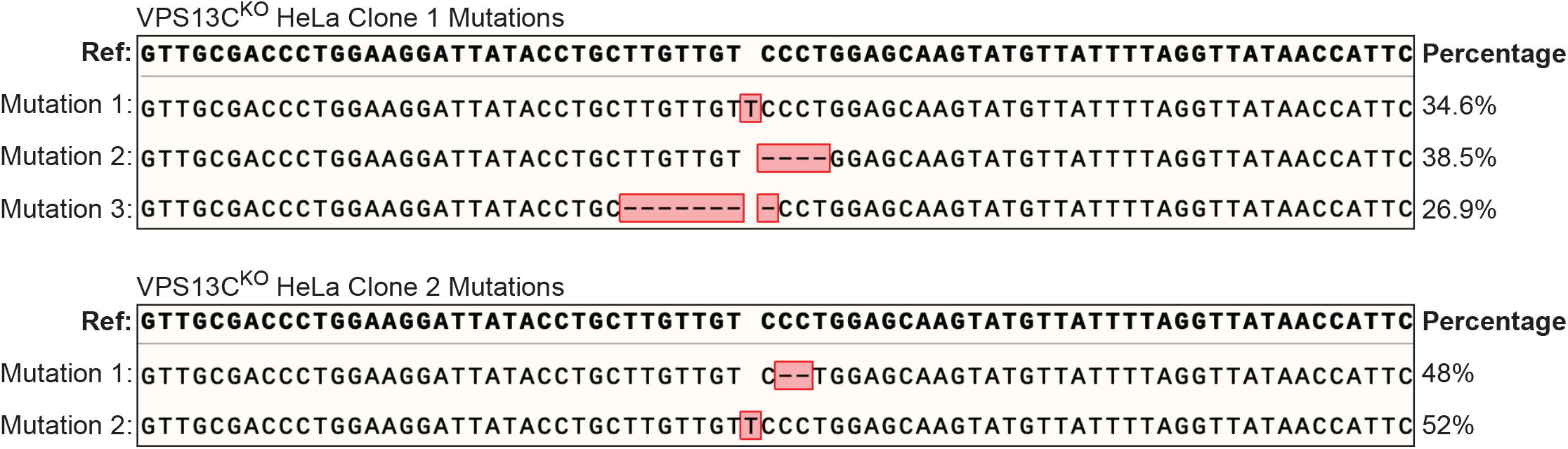
(a) Mutations in the VPS13C locus in VPS13C^KO^ clones 1 and 2. Percent abundance out of 48 bacterial colonies sequenced. The HeLa cells genome is known to be aneuploid.

**Figure S2.**
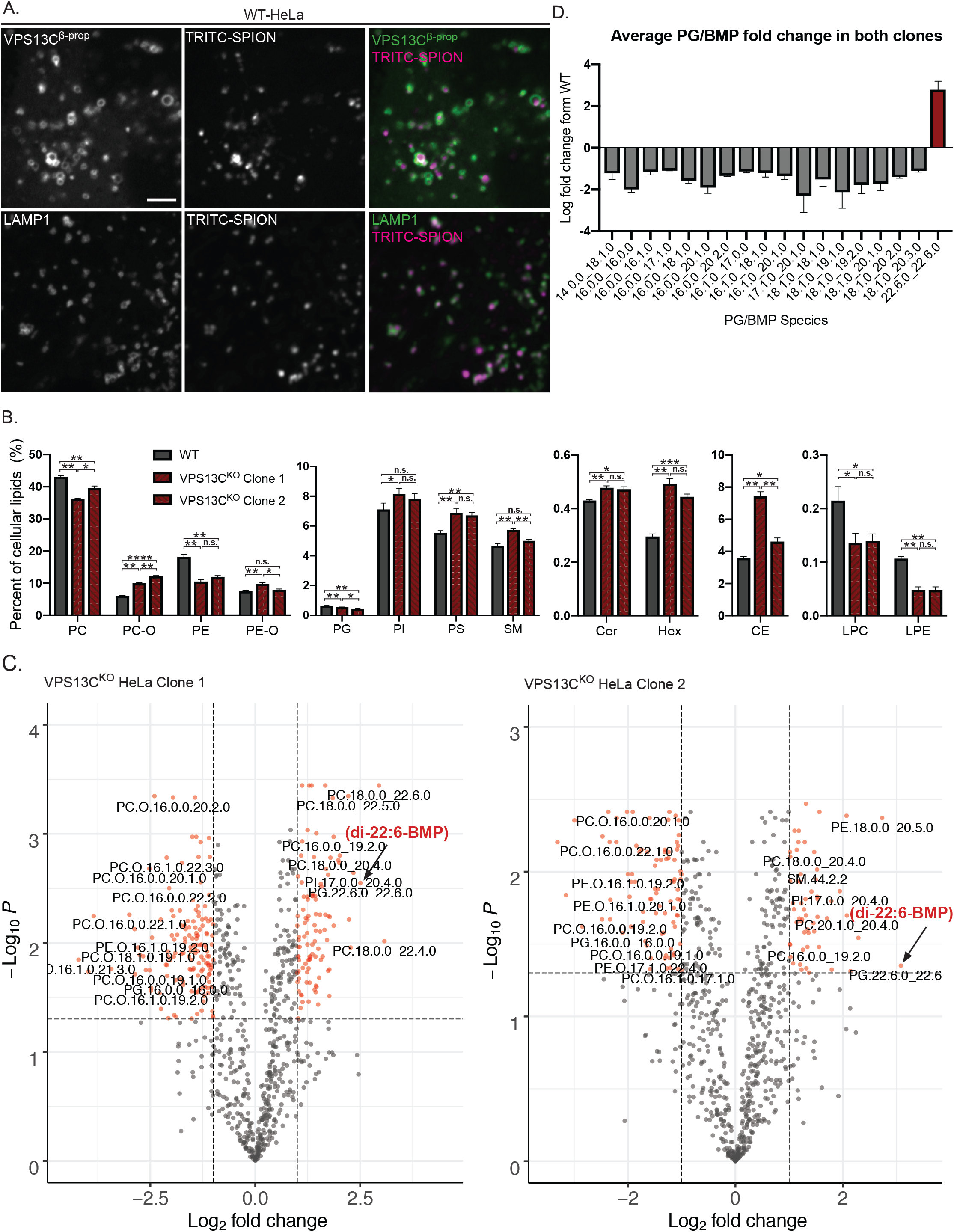
Lipidomics of whole cell and purified lysosomes. (A) Colocalization of TRITC-labelled SPIONs with overexpressed GFP-VPS13C^bprop^ (top row) or LAMP1-GFP (bottom row) after 4 hour pulse and 15 hour chase. Scale bar 5 μm. (B) Percentages of lipid classes in WT and VPS13C^KO^ whole cell lysate normalized to total measured cell lipid content. N = 3 biological replicates. (C) Volcano plots of individual lipid species in VPS13C^KO^ purified lysosomes. Species that surpass the q-value and fold-change thresholds are shown as red dots. Lipid species labels are centered under corresponding dot. Arrows show PG.22.6.0_22.6.0/di-22:6-BMP. (D) Barplot showing the average fold change of PG/BMP species in both clones. Only species with q-values < 0.05 are shown. * q < 0.05, ** q < 0.01, *** q < 0.001, **** q < 0.0001.

**Figure S3.**
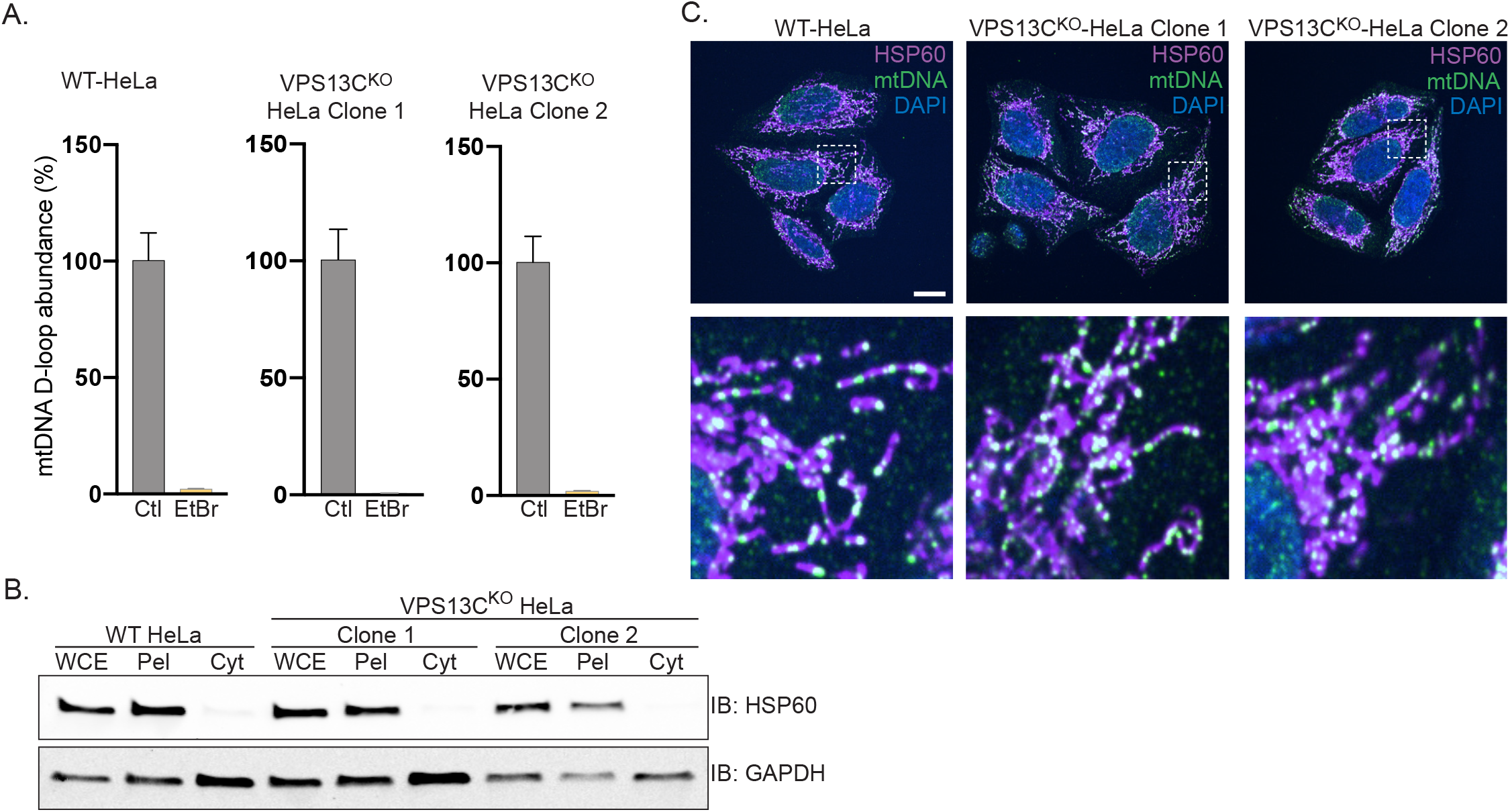
Control experiments for data shown in Figure 4 and normal mtDNA nucleoid morphology in VPS13C^KO^ cells. (A) qPCR of a D-Loop mtDNA amplicon shows efficient depletion of mtDNA after treatment with EtBr. (B) Immunoblot showing that mitochondrial marker HSP60 is present in the whole cell extract (WCE) and pellet (Pel) but absent in the cytosolic fraction (Cyt), while the cytosolic marker GAPDH is present in all fractions. (C) Immunofluorescence showing that mitochondria (magenta) and mtDNA nucleoids (green) have grossly normal morphology in VPS13C^KO^ HeLa cells. Scale bar 20 μm.

**Figure S4.**
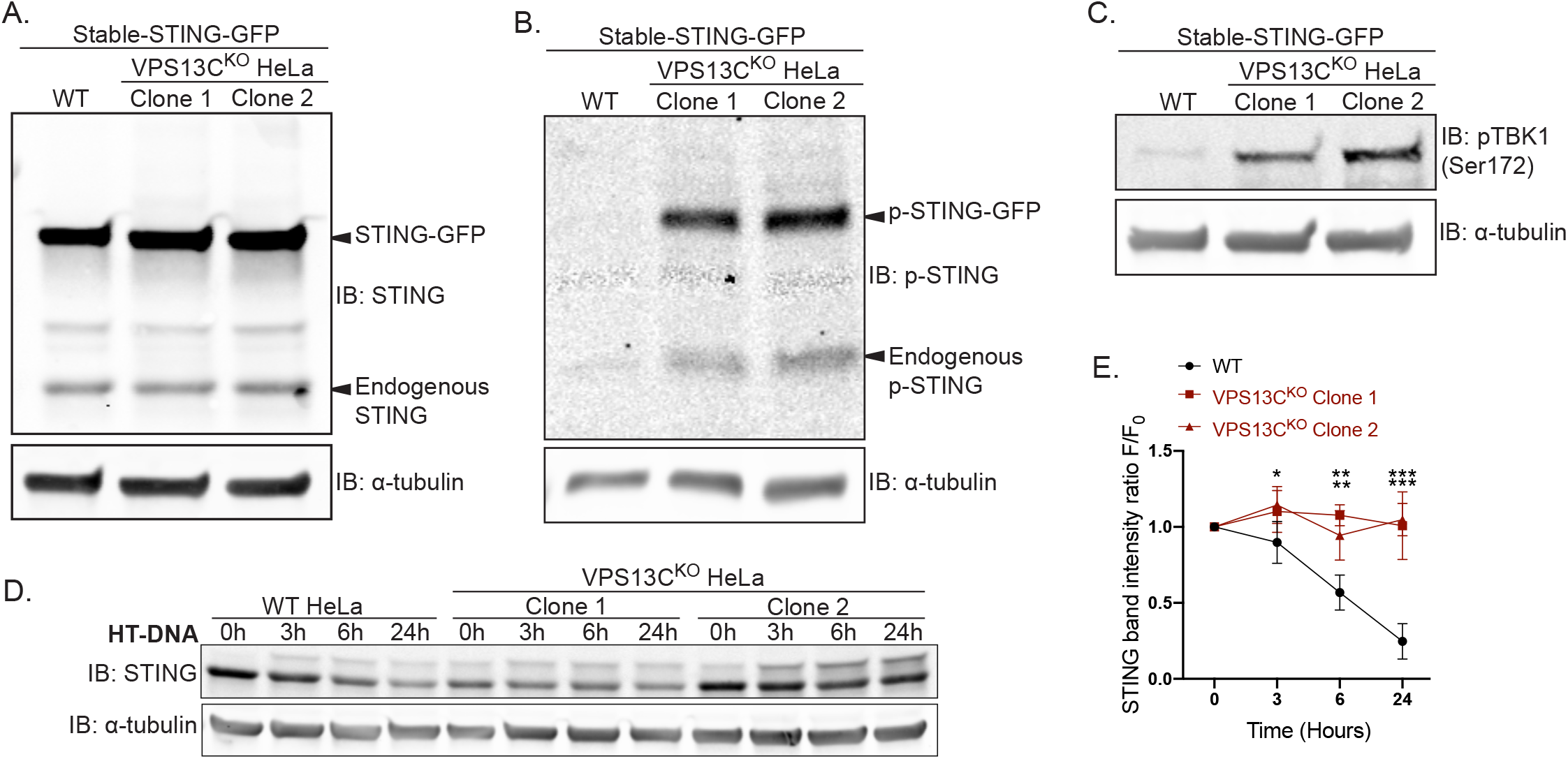
STING signaling is elevated in VPS13C^KO^ cells lines stably expressing STING-GFP. (A) Immunoblot showing similar levels of stable STING-GFP expression in WT and VPS13C^KO^ cells. (B) Immunoblot showing increased levels of p-STING in VPS13C^KO^ cells (both endogenous and STING-GFP). (C) Immunoblot showing increased levels of p-TBK1 in VPS13C^KO^ cells. (D) Timecourse of WT and VPS13C^KO^ cells treated with HT-DNA (500 ng/mL) for 0, 3, 6, and 24 hours. As with cGAMP treatment, STING is not significantly degraded in VPS13C^KO^ cells, quantified in (E). N = 2 biological replicates. * P < 0.05, ** P < 0.01, *** P < 0.001.

**Figure S5.**
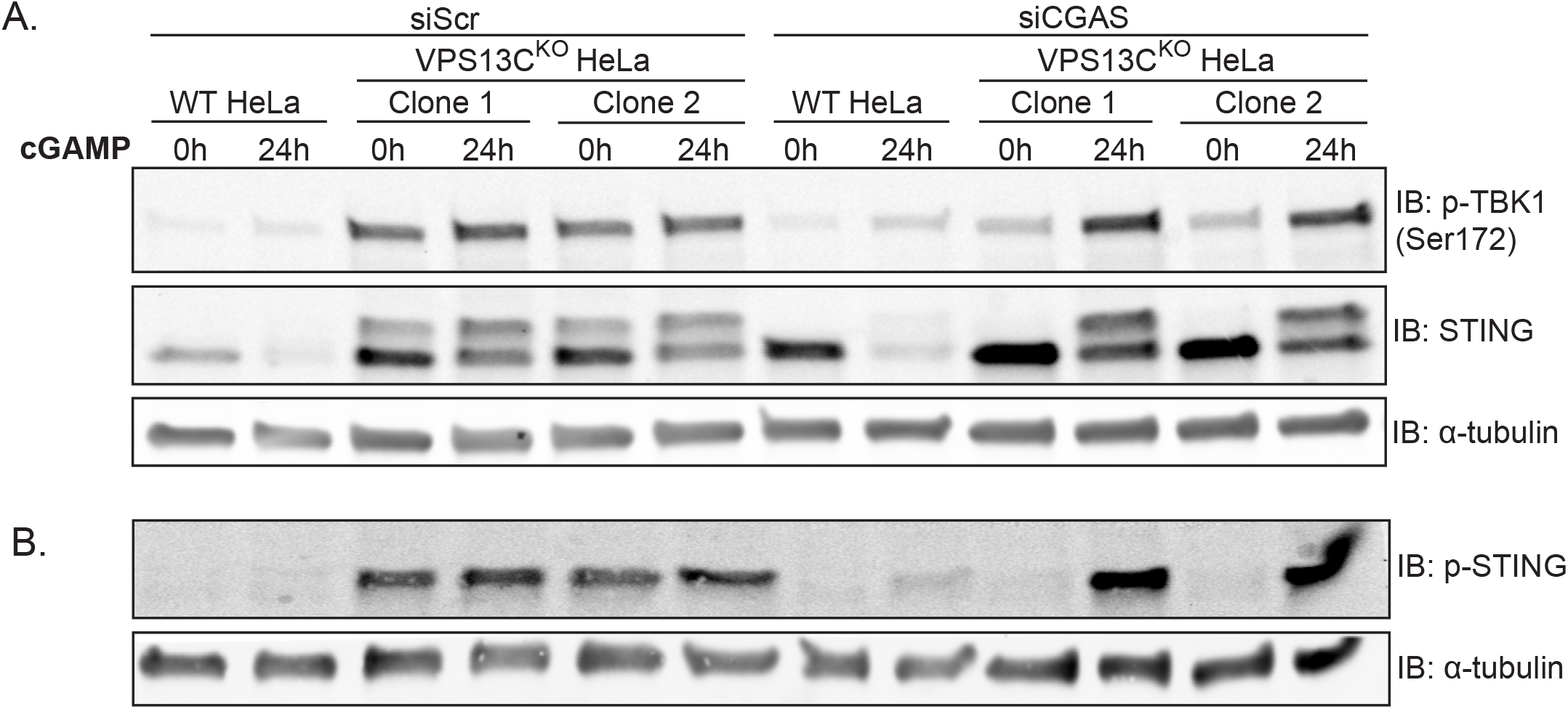
Silencing of cGAS unmasks cGAMP responsiveness and reveals impaired STING degradation in VPS13C^KO^ cells. (A) Intact blot from which the data from Figure 6E and 6G were extracted. (B) Immunoblot of the same samples as in (A) with anti p-STING antibody.

**Figure S6.**
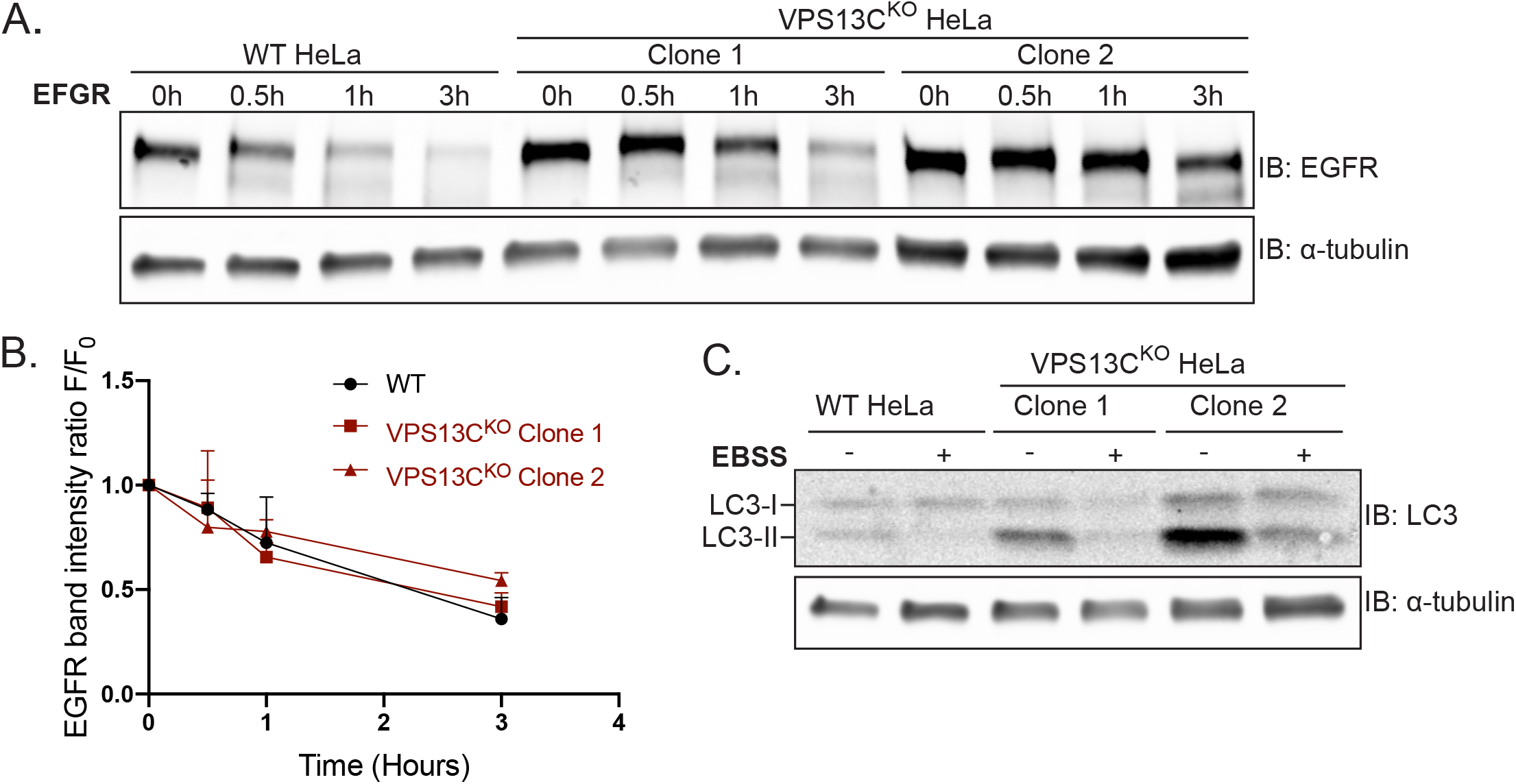
EFGR and LC3 degradation are not impaired in VPS13C^KO^ cells. (A) Immunoblot of EGFR in WT and VPS13C^KO^ cells treated with 100 ng/mL EGF for 0.5, 1, and 3h. (B) Quantification from (A) of band intensity at various timepoints (F) normalized to EGFR intensity at 0h (F0), showing no defect in EGFR degradation kinetics. N = 3 biological replicates. (C) Immunoblot of LC3 in WT and VPS13C^KO^ cells after 6h starvation in Earle’s Balanced Salt Solution (EBSS). Note that the ratio of lipidated LC3-II to LC3-I is elevated under basal conditions in VPS13C^KO^ cells, possibly downstream of STING activation as previously reported (Fischer et al., 2020; Gui et al., 2019).

**Video 1. Treatment with cGAMP induces translocation of STING-GFP from the ER, through the Golgi complex, to dispersed vesicles in WT HeLa cells**. Timelapse of stably expressed STING-GFP in WT HeLa cells after treatment with 50 ug/mL cGAMP just prior to start of imaging, one frame every 15 minutes from 0 to 360 minutes. STING-GFP is localized in an ER-like pattern at 0 min post-treatment, traffics to a Golgi-like pattern 15 min post-treatment, a Golgi/vesicular pattern by 180 minutes post-treatment, and a largely vesicular pattern by 360 min.

## REFERENCES

Abeliovich, A., and A.D. Gitler. 2016. Defects in trafficking bridge Parkinson’s disease pathology and genetics. Nature. 539:207–216.

Alcalay, R.N., F. Hsieh, E. Tengstrand, S. Padmanabhan, M. Baptista, C. Kehoe, S. Narayan, A.K. Boehme, and K. Merchant. 2020. Higher Urine bis(Monoacylglycerol)Phosphate Levels in LRRK2 G2019S Mutation Carriers: Implications for Therapeutic Development. Mov Disord. 35:134–141.

Amick, J., A. Roczniak-Ferguson, and S.M. Ferguson. 2016. C9orf72 binds SMCR8, localizes to lysosomes, and regulates mTORC1 signaling. Mol Biol Cell. 27:3040–3051.

Bonet-Ponce, L., A. Beilina, C.D. Williamson, E. Lindberg, J.H. Kluss, S. Saez-Atienzar, N. Landeck, R. Kumaran, A. Mamais, C.K.E. Bleck, Y. Li, and M.R. Cookson. 2020. LRRK2 mediates tubulation and vesicle sorting from lysosomes. Sci Adv. 6.

Brickner, J.H., and R.S. Fuller. 1997. SOI1 encodes a novel, conserved protein that promotes TGN-endosomal cycling of Kex2p and other membrane proteins by modulating the function of two TGN localization signals. J Cell Biol. 139:23–36.

Campbell, C.L., and P.E. Thorsness. 1998. Escape of mitochondrial DNA to the nucleus in yme1 yeast is mediated by vacuolar-dependent turnover of abnormal mitochondrial compartments. J Cell Sci. 111 (Pt 16):2455–2464.

Cook, D.A., G.T. Kannarkat, A.F. Cintron, L.M. Butkovich, K.B. Fraser, J. Chang, N. Grigoryan, S.A. Factor, A.B. West, J.M. Boss, and M.G. Tansey. 2017. LRRK2 levels in immune cells are increased in Parkinson’s disease. npj Parkinson’s Disease. 3:11.

Darvish, H., P. Bravo, A. Tafakhori, L.J. Azcona, S. Ranji-Burachaloo, A.H. Johari, and C. Paisan-Ruiz. 2018. Identification of a large homozygous VPS13C deletion in a patient with early-onset Parkinsonism. Mov Disord. 33:1968–1970.

Fischer, T.D., C. Wang, B.S. Padman, M. Lazarou, and R.J. Youle. 2020. STING induces LC3B lipidation onto single-membrane vesicles via the V-ATPase and ATG16L1-WD40 domain. J Cell Biol. 219.

Gillingham, A.K., J. Bertram, F. Begum, and S. Munro. 2019. In vivo identification of GTPase interactors by mitochondrial relocalization and proximity biotinylation. Elife. 8.

Gillingham, A.K., R. Sinka, I.L. Torres, K.S. Lilley, and S. Munro. 2014. Toward a comprehensive map of the effectors of rab GTPases. Dev Cell. 31:358–373.

Gkirtzimanaki, K., E. Kabrani, D. Nikoleri, A. Polyzos, A. Blanas, P. Sidiropoulos, A. Makrigiannakis, G. Bertsias, D.T. Boumpas, and P. Verginis. 2018. IFNalpha Impairs Autophagic Degradation of mtDNA Promoting Autoreactivity of SLE Monocytes in a STING-Dependent Fashion. Cell Rep. 25:921–933 e925.

Go, C.D., J.D.R. Knight, A. Rajasekharan, B. Rathod, G.G. Hesketh, K.T. Abe, J.-Y. Youn, P. Samavarchi-Tehrani, H. Zhang, L.Y. Zhu, E. Popiel, J.-P. Lambert, É. Coyaud, S.W.T. Cheung, D. Rajendran, C.J. Wong, H. Antonicka, L. Pelletier, A.F. Palazzo, E.A. Shoubridge, B. Raught, and A.-C. Gingras. 2021. A proximity-dependent biotinylation map of a human cell: an interactive web resource. bioRxiv:796391.

Gonugunta, V.K., T. Sakai, V. Pokatayev, K. Yang, J. Wu, N. Dobbs, and N. Yan. 2017. Trafficking-Mediated STING Degradation Requires Sorting to Acidified Endolysosomes and Can Be Targeted to Enhance Anti-tumor Response. Cell Rep. 21:3234–3242.

Gruenberg, J. 2020. Life in the lumen: The multivesicular endosome. Traffic. 21:76–93.

Gui, X., H. Yang, T. Li, X. Tan, P. Shi, M. Li, F. Du, and Z.J. Chen. 2019. Autophagy induction via STING trafficking is a primordial function of the cGAS pathway. Nature. 567:262–266.

Guillen-Samander, A., X. Bian, and P. De Camilli. 2019. PDZD8 mediates a Rab7-dependent interaction of the ER with late endosomes and lysosomes. Proc Natl Acad Sci U S A. 116:22619–22623.

Guillen-Samander, A., M. Leonzino, M.G. Hanna, N. Tang, H. Shen, and P. De Camilli. 2021. VPS13D bridges the ER to mitochondria and peroxisomes via Miro. J Cell Biol. 220.

Hickman, S., S. Izzy, P. Sen, L. Morsett, and J. El Khoury. 2018. Microglia in neurodegeneration. Nat Neurosci. 21:1359–1369.

Hobert, J.A., and G. Dawson. 2007. A novel role of the Batten disease gene CLN3: association with BMP synthesis. Biochem Biophys Res Commun. 358:111–116.

Hughes, C.E., T.K. Coody, M.Y. Jeong, J.A. Berg, D.R. Winge, and A.L. Hughes. 2020. Cysteine Toxicity Drives Age-Related Mitochondrial Decline by Altering Iron Homeostasis. Cell. 180:296–310 e218.

Khozhukhar, N., D. Spadafora, Y. Rodriguez, and M. Alexeyev. 2018. Elimination of Mitochondrial DNA from Mammalian Cells. Curr Protoc Cell Biol. 78:20 11 21–20 11 14.

Kim, S., Y.C. Wong, F. Gao, and D. Krainc. 2021. Dysregulation of mitochondria-lysosome contacts by GBA1 dysfunction in dopaminergic neuronal models of Parkinson’s disease. Nat Commun. 12:1807.

Kumar, N., M. Leonzino, W. Hancock-Cerutti, F.A. Horenkamp, P. Li, J.A. Lees, H. Wheeler, K.M. Reinisch, and P. De Camilli. 2018. VPS13A and VPS13C are lipid transport proteins differentially localized at ER contact sites. J Cell Biol. 217:3625–3639.

Lang, A.B., A.T. John Peter, P. Walter, and B. Kornmann. 2015. ER-mitochondrial junctions can be bypassed by dominant mutations in the endosomal protein Vps13. J Cell Biol. 210:883–890.

Leonzino, M., K.M. Reinisch, and P. De Camilli. 2021. Insights into VPS13 properties and function reveal a new mechanism of eukaryotic lipid transport. BBA - Molecular and Cell Biology of Lipids.

Lesage, S., V. Drouet, E. Majounie, V. Deramecourt, M. Jacoupy, A. Nicolas, F. Cormier-Dequaire, S.M. Hassoun, C. Pujol, S. Ciura, Z. Erpapazoglou, T. Usenko, C.A. Maurage, M. Sahbatou, S. Liebau, J. Ding, B. Bilgic, M. Emre, N. Erginel-Unaltuna, G. Guven, F. Tison, C. Tranchant, M. Vidailhet, J.C. Corvol, P. Krack, A.L. Leutenegger, M.A. Nalls, D.G. Hernandez, P. Heutink, J.R. Gibbs, J. Hardy, N.W. Wood, T. Gasser, A. Durr, J.F. Deleuze, M. Tazir, A. Destee, E. Lohmann, E. Kabashi, A. Singleton, O. Corti, A. Brice, S. French Parkinson’s Disease Genetics, and C. International Parkinson’s Disease Genomics. 2016. Loss of VPS13C Function in Autosomal-Recessive Parkinsonism Causes Mitochondrial Dysfunction and Increases PINK1/Parkin-Dependent Mitophagy. Am J Hum Genet. 98:500–513.

Li, P., J.A. Lees, C.P. Lusk, and K.M. Reinisch. 2020. Cryo-EM reconstruction of a VPS13 fragment reveals a long groove to channel lipids between membranes. J Cell Biol. 219.

Lin, R., C. Heylbroeck, P.M. Pitha, and J. Hiscott. 1998. Virus-dependent phosphorylation of the IRF-3 transcription factor regulates nuclear translocation, transactivation potential, and proteasome-mediated degradation. Mol Cell Biol. 18:2986–2996.

Liu, N., E.A. Tengstrand, L. Chourb, and F.Y. Hsieh. 2014. Di-22:6-bis(monoacylglycerol)phosphate: A clinical biomarker of drug-induced phospholipidosis for drug development and safety assessment. Toxicol Appl Pharmacol. 279:467–476.

Liu, S., X. Cai, J. Wu, Q. Cong, X. Chen, T. Li, F. Du, J. Ren, Y.T. Wu, N.V. Grishin, and Z.J. Chen. 2015. Phosphorylation of innate immune adaptor proteins MAVS, STING, and TRIF induces IRF3 activation. Science. 347:aaa2630.

Liu, X., K. Salokas, F. Tamene, Y. Jiu, R.G. Weldatsadik, T. Ohman, and M. Varjosalo. 2018. An AP-MS- and BioID-compatible MAC-tag enables comprehensive mapping of protein interactions and subcellular localizations. Nat Commun. 9:1188.

Logan, T., S. DeVos, M.J. Simon, S. Davis, J. Wang, R. Low, F. Huang, Y. Rajendra, R. Prorok, E. Sun, A. Rana, J. Hsiao-Nakamoto, S. Mosesova, Y. Zhu, G.M. Cherf, B. Lengerich, A. Bhalla, D.J. Kim, D. Chan, J. Duque, H. Tran, M. Lenser, H. Nguyen, R. Chau, T. Earr, M.S. Kariolis, K.M. Monroe, P.E. Sanchez, M.S. Dennis, K.R. Henne, K. Gunasekaran, G. Astarita, R.J. Watts, Z.K. Sweeney, J.W. Lewcock, A. Srivastava, and G. Di Paolo. 2020. A brain penetrant progranulin biotherapeutic rescues lysosomal and inflammatory phenotypes in the brain of GRN knockout mice. Alzheimer’s & Dementia. 16:e040602.

Malpartida, A.B., M. Williamson, D.P. Narendra, R. Wade-Martins, and B.J. Ryan. 2021. Mitochondrial Dysfunction and Mitophagy in Parkinson’s Disease: From Mechanism to Therapy. Trends Biochem Sci. 46:329–343.

McCauley, M.E., J.G. O’Rourke, A. Yanez, J.L. Markman, R. Ho, X. Wang, S. Chen, D. Lall, M. Jin, A. Muhammad, S. Bell, J. Landeros, V. Valencia, M. Harms, M. Arditi, C. Jefferies, and R.H. Baloh. 2020. C9orf72 in myeloid cells suppresses STING-induced inflammation. Nature. 585:96–101.

Motwani, M., S. Pesiridis, and K.A. Fitzgerald. 2019. DNA sensing by the cGAS-STING pathway in health and disease. Nat Rev Genet. 20:657–674.

Nalls, M.A., C. Blauwendraat, C.L. Vallerga, K. Heilbron, S. Bandres-Ciga, D. Chang, M. Tan, D.A. Kia, A.J. Noyce, A. Xue, J. Bras, E. Young, R. von Coelln, J. Simon-Sanchez, C. Schulte, M. Sharma, L. Krohn, L. Pihlstrom, A. Siitonen, H. Iwaki, H. Leonard, F. Faghri, J.R. Gibbs, D.G. Hernandez, S.W. Scholz, J.A. Botia, M. Martinez, J.C. Corvol, S. Lesage, J. Jankovic, L.M. Shulman, M. Sutherland, P. Tienari, K. Majamaa, M. Toft, O.A. Andreassen, T. Bangale, A. Brice, J. Yang, Z. Gan-Or, T. Gasser, P. Heutink, J.M. Shulman, N.W. Wood, D.A. Hinds, J.A. Hardy, H.R. Morris, J. Gratten, P.M. Visscher, R.R. Graham, A.B. Singleton T. and Me Research, C. System Genomics of Parkinson’s Disease, and C. International Parkinson’s Disease Genomics. 2019. Identification of novel risk loci, causal insights, and heritable risk for Parkinson’s disease: a meta-analysis of genome-wide association studies. Lancet Neurol. 18:1091–1102.

Oka, T., S. Hikoso, O. Yamaguchi, M. Taneike, T. Takeda, T. Tamai, J. Oyabu, T. Murakawa, H. Nakayama, K. Nishida, S. Akira, A. Yamamoto, I. Komuro, and K. Otsu. 2012. Mitochondrial DNA that escapes from autophagy causes inflammation and heart failure. Nature. 485:251–255.

Okamoto, S., Y. Amaishi, I. Maki, T. Enoki, and J. Mineno. 2019. Highly efficient genome editing for single-base substitutions using optimized ssODNs with Cas9-RNPs. Sci Rep. 9:4811.

Park, J.S., M.K. Thorsness, R. Policastro, L.L. McGoldrick, N.M. Hollingsworth, P.E. Thorsness, and A.M. Neiman. 2016. Yeast Vps13 promotes mitochondrial function and is localized at membrane contact sites. Mol Biol Cell. 27:2435–2449.

Pickrell, A.M., and R.J. Youle. 2015. The roles of PINK1, parkin, and mitochondrial fidelity in Parkinson’s disease. Neuron. 85:257–273.

Ran, F.A., P.D. Hsu, J. Wright, V. Agarwala, D.A. Scott, and F. Zhang. 2013. Genome engineering using the CRISPR-Cas9 system. Nat Protoc. 8:2281–2308.

Riley, J.S., and S.W. Tait. 2020. Mitochondrial DNA in inflammation and immunity. EMBO Rep. 21:e49799.

Saunders, A., E.Z. Macosko, A. Wysoker, M. Goldman, F.M. Krienen, H. de Rivera, E. Bien, M. Baum, L. Bortolin, S. Wang, A. Goeva, J. Nemesh, N. Kamitaki, S. Brumbaugh, D. Kulp, and S.A. McCarroll. 2018. Molecular Diversity and Specializations among the Cells of the Adult Mouse Brain. Cell. 174:1015–1030 e1016.

Schormair, B., D. Kemlink, B. Mollenhauer, O. Fiala, G. Machetanz, J. Roth, R. Berutti, T.M. Strom, B. Haslinger, C. Trenkwalder, D. Zahorakova, P. Martasek, E. Ruzicka, and J. Winkelmann. 2018. Diagnostic exome sequencing in early-onset Parkinson’s disease confirms VPS13C as a rare cause of autosomal-recessive Parkinson’s disease. Clin Genet. 93:603–612.

Shang, G., C. Zhang, Z.J. Chen, X.C. Bai, and X. Zhang. 2019. Cryo-EM structures of STING reveal its mechanism of activation by cyclic GMP-AMP. Nature. 567:389–393.

Showalter, M.R., A.L. Berg, A. Nagourney, H. Heil, K.L. Carraway, 3rd, and O. Fiehn. 2020. The Emerging and Diverse Roles of Bis(monoacylglycero) Phosphate Lipids in Cellular Physiology and Disease. Int J Mol Sci. 21.

Sliter, D.A., J. Martinez, L. Hao, X. Chen, N. Sun, T.D. Fischer, J.L. Burman, Y. Li, Z. Zhang, D.P. Narendra, H. Cai, M. Borsche, C. Klein, and R.J. Youle. 2018. Parkin and PINK1 mitigate STING-induced inflammation. Nature. 561:258–262.

Smolders, S., S. Philtjens, D. Crosiers, A. Sieben, E. Hens, B. Heeman, S. Van Mossevelde, P. Pals, B. Asselbergh, R. Dos Santos Dias, Y. Vermeiren, R. Vandenberghe, S. Engelborghs, P.P. De Deyn, J.J. Martin, P. Cras, W. Annaert, C. Van Broeckhoven, and B. consortium. 2021. Contribution of rare homozygous and compound heterozygous VPS13C missense mutations to dementia with Lewy bodies and Parkinson’s disease. Acta Neuropathol Commun. 9:25.

Sprenger, H.G., T. MacVicar, A. Bahat, K.U. Fiedler, S. Hermans, D. Ehrentraut, K. Ried, D. Milenkovic, N. Bonekamp, N.G. Larsson, H. Nolte, P. Giavalisco, and T. Langer. 2021. Cellular pyrimidine imbalance triggers mitochondrial DNA-dependent innate immunity. Nat Metab. 3:636–650.

Swanson, K.V., R.D. Junkins, C.J. Kurkjian, E. Holley-Guthrie, A.A. Pendse, R. El Morabiti, A. Petrucelli, G.N. Barber, C.A. Benedict, and J.P. Ting. 2017. A noncanonical function of cGAMP in inflammasome priming and activation. J Exp Med. 214:3611–3626.

Tharkeshwar, A.K., D. Demedts, and W. Annaert. 2020. Superparamagnetic Nanoparticles for Lysosome Isolation to Identify Spatial Alterations in Lysosomal Protein and Lipid Composition. STAR Protoc. 1:100122.

Tharkeshwar, A.K., J. Trekker, W. Vermeire, J. Pauwels, R. Sannerud, D.A. Priestman, D. Te Vruchte, K. Vints, P. Baatsen, J.P. Decuypere, H. Lu, S. Martin, P. Vangheluwe, J.V. Swinnen, L. Lagae, F. Impens, F.M. Platt, K. Gevaert, and W. Annaert. 2017. A novel approach to analyze lysosomal dysfunctions through subcellular proteomics and lipidomics: the case of NPC1 deficiency. Sci Rep. 7:41408.

Thorsness, P.E., and T.D. Fox. 1993. Nuclear mutations in Saccharomyces cerevisiae that affect the escape of DNA from mitochondria to the nucleus. Genetics. 134:21–28.

Ugur, B., W. Hancock-Cerutti, M. Leonzino, and P. De Camilli. 2020. Role of VPS13, a protein with similarity to ATG2, in physiology and disease. Curr Opin Genet Dev. 65:61–68.

Vidyadhara, D.J., J.E. Lee, and S.S. Chandra. 2019. Role of the endolysosomal system in Parkinson’s disease. J Neurochem. 150:487–506.

Weindel, C.G., S.L. Bell, K.J. Vail, K.O. West, K.L. Patrick, and R.O. Watson. 2020. LRRK2 maintains mitochondrial homeostasis and regulates innate immune responses to Mycobacterium tuberculosis. Elife. 9.

West, A.B., D.J. Moore, S. Biskup, A. Bugayenko, W.W. Smith, C.A. Ross, V.L. Dawson, and T.M. Dawson. 2005. Parkinson’s disease-associated mutations in leucine-rich repeat kinase 2 augment kinase activity. Proc Natl Acad Sci U S A. 102:16842–16847.

West, A.P., W. Khoury-Hanold, M. Staron, M.C. Tal, C.M. Pineda, S.M. Lang, M. Bestwick, B.A. Duguay, N. Raimundo, D.A. MacDuff, S.M. Kaech, J.R. Smiley, R.E. Means, A. Iwasaki, and G.S. Shadel. 2015. Mitochondrial DNA stress primes the antiviral innate immune response. Nature. 520:553–557.

West, A.P., and G.S. Shadel. 2017. Mitochondrial DNA in innate immune responses and inflammatory pathology. Nat Rev Immunol. 17:363–375.

Wong, Y.C., D. Ysselstein, and D. Krainc. 2018. Mitochondria-lysosome contacts regulate mitochondrial fission via RAB7 GTP hydrolysis. Nature. 554:382–386.

Yambire, K.F., C. Rostosky, T. Watanabe, D. Pacheu-Grau, S. Torres-Odio, A. Sanchez-Guerrero, O. Senderovich, E.G. Meyron-Holtz, I. Milosevic, J. Frahm, A.P. West, and N. Raimundo. 2019. Impaired lysosomal acidification triggers iron deficiency and inflammation in vivo. Elife. 8.

Yeshaw, W.M., M. van der Zwaag, F. Pinto, L.L. Lahaye, A.I. Faber, R. Gomez-Sanchez, A.M. Dolga, C. Poland, A.P. Monaco, I.S.C. van, N.A. Grzeschik, A. Velayos-Baeza, and O.C. Sibon. 2019. Human VPS13A is associated with multiple organelles and influences mitochondrial morphology and lipid droplet motility. Elife. 8.

Yu, C.H., S. Davidson, C.R. Harapas, J.B. Hilton, M.J. Mlodzianoski, P. Laohamonthonkul, C. Louis, R.R.J. Low, J. Moecking, D. De Nardo, K.R. Balka, D.J. Calleja, F. Moghaddas, E. Ni, C.A. McLean, A.L. Samson, S. Tyebji, C.J. Tonkin, C.R. Bye, B.J. Turner, G. Pepin, M.P. Gantier, K.L. Rogers, K. McArthur, P.J. Crouch, and S.L. Masters. 2020. TDP-43 Triggers Mitochondrial DNA Release via mPTP to Activate cGAS/STING in ALS. Cell. 183:636–649 e618.

Zhang, C., G. Shang, X. Gui, X. Zhang, X.C. Bai, and Z.J. Chen. 2019. Structural basis of STING binding with and phosphorylation by TBK1. Nature. 567:394–398.

