## Supplemental Table 1 for "ER-lysosome lipid transfer protein VPS13C/PARK23 prevents aberrant mtDNA-dependent STING signaling"

| Table S1: List of oligos used in this study | |  |  |
| --- | --- | --- | --- |
| **CRISPR oligos** |  |  |  |
| **Oligo** | **Cloning Method** | **Sequence** | **Backbone** |
| VPS13C KO gRNA | T7 Ligase | CACCGTTATACCTGCTTGTTGTCCC | PX459 |
|  |  | AAACGGGACAACAAGCAGGTATAAC |  |
| VPS13C Repair gRNA | T7 Ligase | CACCGTATACCTGCTTGTTGTTCCC | PX459 |
|  |  | AAACGGGAACAACAAGCAGGTATAAC |  |
| **Oligo** | **Sequence** |  |  |
| VPS13C ssODN Repair template | TGGTTATAACCTAAAATAACATACTTGCTCCAGGGACAACAAGCAGGTATAATCCTTCCAGGGTCGCAACA | |  |
| VPS13C-CRISPR-Sequencing-F-Xho1 | GCACACTCGAGCTGAGTGATGTACATACCATGAAGAG    GCACAGGGCCCCACAAATGCGGTAAGATATATGTGC | |  |
| VPS13C-CRISPR-Sequencing-R-Apa1 |  |  |  |
| **siRNA** |  |  |  |
| **Target** | **Company** | **siRNA ID** |  |
| CGAS (MB21D1) | ThermoFisher | s41746 |  |
| STING (TMEM173) | ThermoFisher | s50644 |  |
| **qPCR primers** |  |  |  |
| **Target** | **Sequence** | |  |
| hβ-Actin-F | CCTGGCACCCAGCACAAT | |  |
| hβ-Actin-R | GCCGATCCACACGGAGTACT | |  |
| hIFIT1‐F | TTGATGACGATGAAATGCCTGA | |  |
| hIFIT1‐R | CAGGTCACCAGACTCCTCAC | |  |
| hIFIT3‐F | TCAGAAGTCTAGTCACTTGGGG | |  |
| hIFIT3‐R | ACACCTTCGCCCTTTCATTTC | |  |
| hOasl-F | CTGATGCAGGAACTGTATAGCAC | |  |
| hOasl-R | CACAGCGTCTAGCACCTCTT | |  |
| hSTAT1-F | CAGCTTGACTCAAAATTCCTGGA | |  |
| hSTAT1-R | TGAAGATTACGCTTGCTTTTCCT | |  |
| hISG15-F | CGCAGATCACCCAGAAGATCG | |  |
| hISG15-R | TTCGTCGCATTTGTCCACCA | |  |
| hB2M-F | TGCTGTCTCCATGTTTGATGTATCT | |  |
| hB2M-R | TCTCTGCTCCCCACCTCTAAGT | |  |
| h-mt-Dloop-F | CCGTGAGTGGTTAATAGGGTGATA | |  |
| h-mt-Dloop-R | CATAAAGCCTAAATAGCCCACACG | |  |
| h-mt-CYB-F | CCATCCTCCATATATCCAAA | |  |
| h-mt-CYB-R | CCAATGATGGTAAAAGGGTA | |  |
| h-mt-ND4-F | CCTCGTAGTAACAGCCATTC | |  |
| h-mt-ND4-R | TTGAAGTCCTTGAGAGAGGA | |  |
